# *De novo* Design of a Vanadium Haloperoxidase

**DOI:** 10.64898/2026.10.16.759729

**Authors:** Ryan A. Herold, Lancia A. Carson, Kevin J. Anderton, Kent J. Griffith, F. Akif Tezcan

## Abstract

Organohalogens are an essential class of compounds for both synthetic chemistry and natural systems, yet the design of halogenases to catalyze carbon–halogen (C–X) bond formation has remained elusive. Vanadium-dependent haloperoxidases (V-HPOs) are attractive targets for such an effort, but they pose an unusual design problem: the vanadate (VO_4_^3−^) cofactor is an anionic oxometalate that must be held by a precisely arranged network of hydrogen-bonding and electrostatic interactions rather than by the strong coordinate-covalent bonds through which proteins bind cationic transition metals. We addressed this challenge by utilizing a machine-learning-guided workflow to design VAN1, a highly stable 20 kDa protein that binds VO_4_^3−^ and brominates activated aromatic substrates, including *L*-tyrosine. X-ray crystal structures of metal-bound VAN1 closely matched the computational model and confirmed the designed active site, and structure-guided mutagenesis improved VO_4_^3−^ affinity 15-fold and bromination activity 2.6-fold, in quantitative agreement with the resulting change in cofactor occupancy. VAN1 remains folded in neat organic solvents and is substantially more active in aqueous/organic mixtures than in water, representing an important advantage for synthetic applications. As the first example of a *de novo* enzyme that catalyzes C–X bond formation, VAN1 establishes a new starting point for V-HPO engineering and mechanistic studies.

## Main Text

Halogenated organic compounds are essential for modern chemistry, serving as important synthetic intermediates and increasing the bioactivities of myriad pharmaceuticals and agrochemicals.^1, 2, 3^ Likewise, organohalogens play crucial roles in natural systems (e.g., hormones, antibiotics, halogenated aerosols)^4, 5^ and thus organisms have evolved five types of halogenases which can catalyze the challenging formation of C–X bonds under mild aqueous conditions.^6, 7^ Among these enzymes, vanadium-dependent haloperoxidases (V-HPOs) stand out for their potential use in industrial settings due to their redox stability, broad substrate scope, and lack of required expensive cofactors/cosubstrates or auxiliary enzymes.^8^ Despite significant interest in engineering/evolving natural enzymes for halogenation chemistry, the *de novo* design of a halogenase enzyme has thus far remained out of reach. To address this gap, we chose to design a V-dependent haloperoxidase to extend the field of protein design to this highly important reaction class.

Vanadium is the second most abundant transition metal in the ocean,^9, 10^ yet V-HPOs represent one of only two known V-dependent enzymes, with the other being V nitrogenase. Furthermore, despite being present in evolutionarily diverse organisms including algae, fungi, and bacteria, only one highly conserved active site architecture has evolved to facilitate V-catalyzed haloperoxidation (Fig. S1).^11, 12^ Thus, only two protein folds have evolved to utilize V as a catalyst despite it being redox active, abundant in soluble form, and capable of catalyzing a wide variety of chemical transformations.^13, 14^ The notable lack of V-dependent enzymes may reflect the chemistry of solvated V species, which primarily exist as oxometalates (e.g., VO_4_^3−^). In contrast to other first-row transition metals (e.g., Fe, Cu, Zn) which are generally cationic, have labile ligands, and form strong coordinate-covalent bonds with protein side chains, vanadate is anionic and maintains strong V–O bonds which limit direct protein–metal interactions (Fig. 1a). Thus, the binding of V by proteins is governed by entirely different biochemical rules driven by precise electrostatic and hydrogen-bonding networks (Fig. 1a), posing a formidable challenge for computational protein design.

**Fig. 1.**
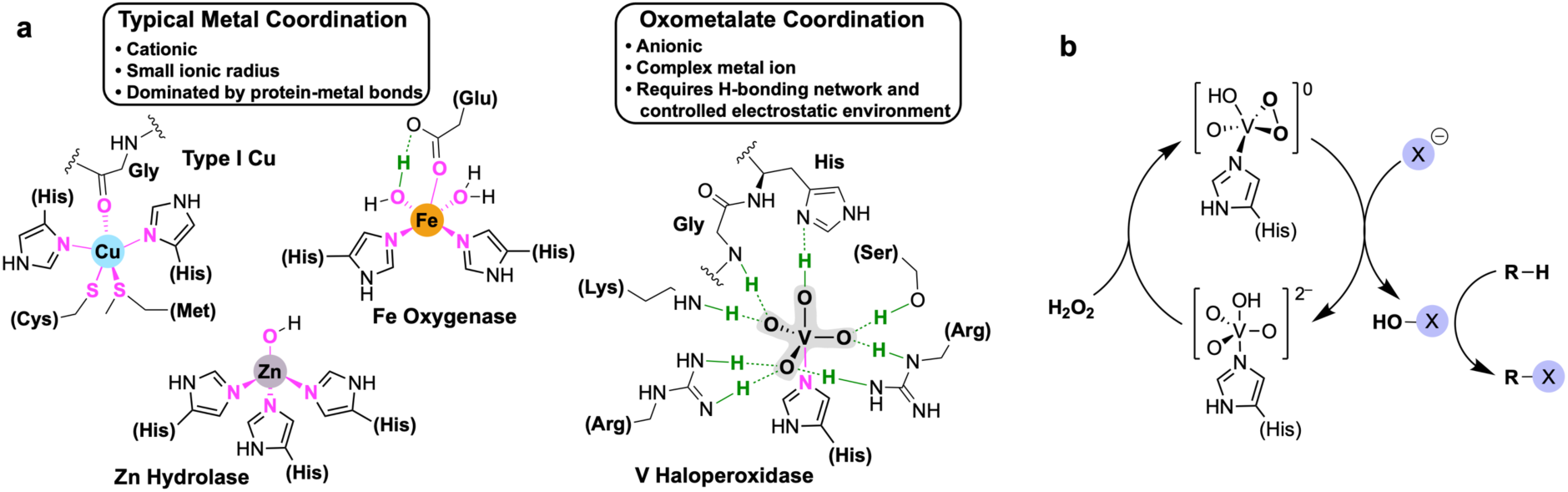
Typical first-row transition metal versus oxometalate protein binding and the V-HPO catalytic cycle. (a) (left) Example metal binding sites for first-row transition metals commonly found in biology: type I Cu (PDB: 4AZU),^27^ Fe oxygenase (PDB: 1ZZ9),^28^ Zn hydrolase (PDB: 2CLT).^29^ (right) Vanadate bound by a V-HPO showing the single V-coordinating His and the stabilizing H-bonding network (PDB: 1IDQ).^15^ Metal-ligand bonds are colored magenta, and hydrogen bonds are colored green. (b) V-HPO mechanism in which the halogenating species is diffusible HOX which can react with substrate at/near the active site for selective halogenation or in solution for non-selective halogenation.

Natural V-HPOs bind the VO_4_^3−^ cofactor through a single His-V bond supported by a shell of hydrogen bonds provided by conserved His, Arg, Lys, Ser and Gly residues (Fig. 1a).^15, 16, 17^ The VO_4_^3−^ center reacts with H_2_O_2_ to form an activated, side-on V-peroxo intermediate that oxidizes halides (Cl^−^, Br^−^, or I^−^) to yield electrophilic halogenating species that may react with substrates at/near the active site for selective halogenation^18^ or diffuse away from the enzyme for non-selective chemistry (Fig. 1b).^2, 19^ Because nature offers only a single structural template for this chemistry, bioengineering efforts have been inherently constrained. In fact, the only example of an artificial V-HPO is a structurally homologous acid phosphatase which has an identical conserved primary coordination sphere to natural V-HPOs.^11, 20, 21^ Thus, its ability to bind VO_4_^3−^ (in place of PO_4_^3−^) and mimic V-HPO activity is unsurprising.^1^ Synthetic V^V^ complexes can also mimic haloperoxidase activity, however they exhibit significantly lower turnover rates compared to the natural enzymes,^14, 22, 23^ underscoring the importance of the highly organized secondary coordination sphere of natural V-HPOs. Thus, to exploit the evolvability and tunability of enzymes as three-dimensional ligand scaffolds, and to provide a new starting point for mechanistic studies, we utilized a machine-learning-guided protein design workflow, which has proven successful for the design of completely new protein folds around targeted structural templates,^24, 25, 26^ to design a *de novo* V-HPO.

## Results and Discussion

### Protein Design Approach

To design a protein capable of binding and activating VO_4_^3−^ for V-HPO catalysis, we selected six active-site residues from the model *C. inaequalis* chloroperoxidase^15^ to form the starting motif (Fig. 2). This motif included the V-coordinating histidine, four positively charged residues that stabilize VO_4_^3−^ through H-bonding interactions, and one secondary-sphere Asp that forms a salt bridge with a conserved arginine (Fig. 2). We used RFdiffusion^30^ to generate protein backbones in iterative batches, applying a guiding potential to prevent protein ingression into the VO_4_^3−^ binding pocket. Importantly, because we conserved only six residues from the natural enzyme, the designed protein folds had essentially no resemblance to the native chloroperoxidase. However, stabilizing six conserved residues, each oriented in different positions around the VO_4_^3−^ center, proved challenging and initial designs frequently placed these conserved residues on flexible loops that would be unlikely to adopt the desired geometry in solution. To overcome this, we first screened for designs which anchored the conserved residues on rigid secondary structures (α-helices or β-sheets). We then aligned each backbone with the starting motif to ensure that the VO_4_^3−^ pocket remained vacant, discarding designs where the generated backbone occluded the active site. Finally, we selected backbones featuring a well-defined, solvent-exposed VO_4_^3−^ binding pocket that would allow small-molecule substrate access to the VO_4_^3−^ center.

**Fig. 2.**
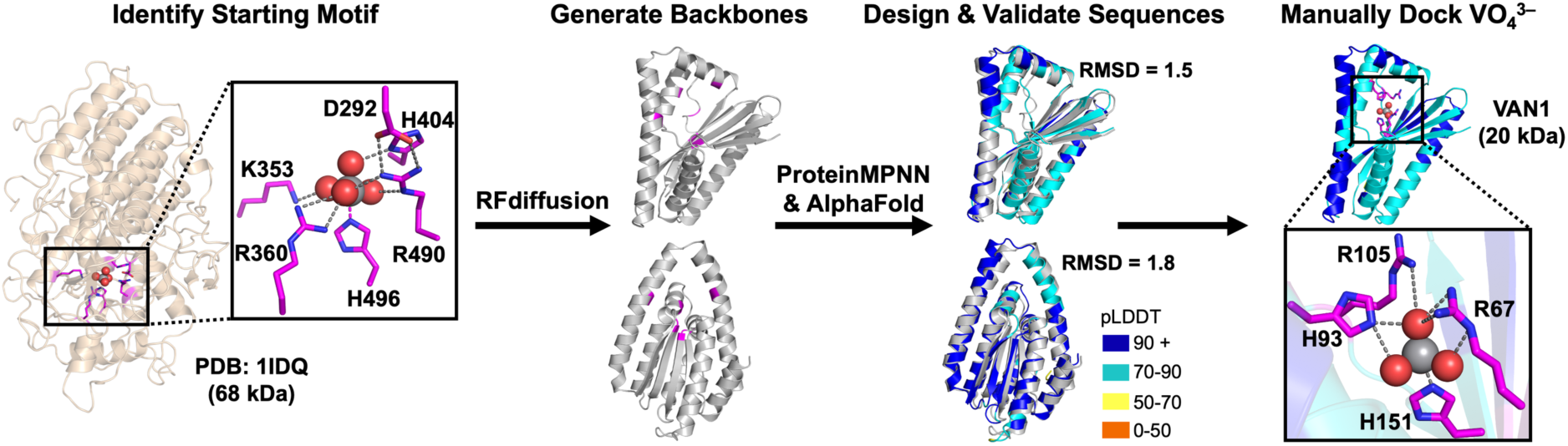
Machine-learning-guided design approach used in this work.

Using ProteinMPNN,^31^ we designed sequences for the best candidate scaffolds (Fig. 2). Cysteine and additional histidine residues were excluded to prevent disulfide bond formation and non-specific metal binding. AlphaFold2^32^ was used to predict three-dimensional structures, and we used pLDDT scores and RMSD values obtained from alignment with the RFdiffusion backbones as the primary scoring metrics. As a final *in silico* validation step, we manually docked VO_4_^3−^ into the binding pockets of top-scoring AlphaFold2 predicted structures to check for possible steric clashes and confirm that the positionally-conserved His and Arg residues could potentially coordinate V and provide stabilizing H-bonding interactions (Fig. 2). Of >1000 backbones generated using different input parameters, only three scaffolds passed our screening criteria, from which we selected nine protein sequences for expression and characterization (Fig. S2).

### Bromoperoxidase Activity of VAN1

All nine sequences were expressed in *E. coli*, however only three of the proteins were sufficiently soluble to be isolated in high yields (Fig. S2). The low solubility may reflect the highly unusual protein folds that were required to accommodate the six conserved residues while maintaining stable secondary structures within a sequence length of only 180–240 residues. The soluble proteins, VAN1, VAN6 and VAN7, were assayed for bromoperoxidase activity using the well-established monochlorodimedone (MCD) assay (Fig. 3a).^33, 34^ Neither VAN6 nor VAN7 showed bromoperoxidase activity above the low background rate catalyzed by VO_4_^3−^ in solution. Excitingly, VAN1 exhibited rates of bromination that were 5–9-fold higher than the background over a period of 4 h (Fig. 3b). Bromination rates increased linearly with the amount of VAN1 added (Fig. S3i), and the rate dependence on [Br^−^] approached saturation, consistent with Michaelis–Menten kinetics (Fig. 3c). The *K*_m_ for Br^−^ was 550 mM. While this value indicates a much lower affinity for Br^−^ compared to natural V-BPOs (*K*_m_ = 2–18 mM),^4, 12, 35^ the VAN1 affinity is comparable to bacterial acid phosphatases which have an identical binding site to natural V-HPOs (*K*_m_ = 100–200 mM).^35^ Thus, despite VAN1 possessing only four positionally-conserved primary-sphere residues from the natural enzyme, these experiments confirm that our minimal binding site possesses functional, baseline activity within a *de novo* scaffold prior to any optimization.

**Fig. 3.**
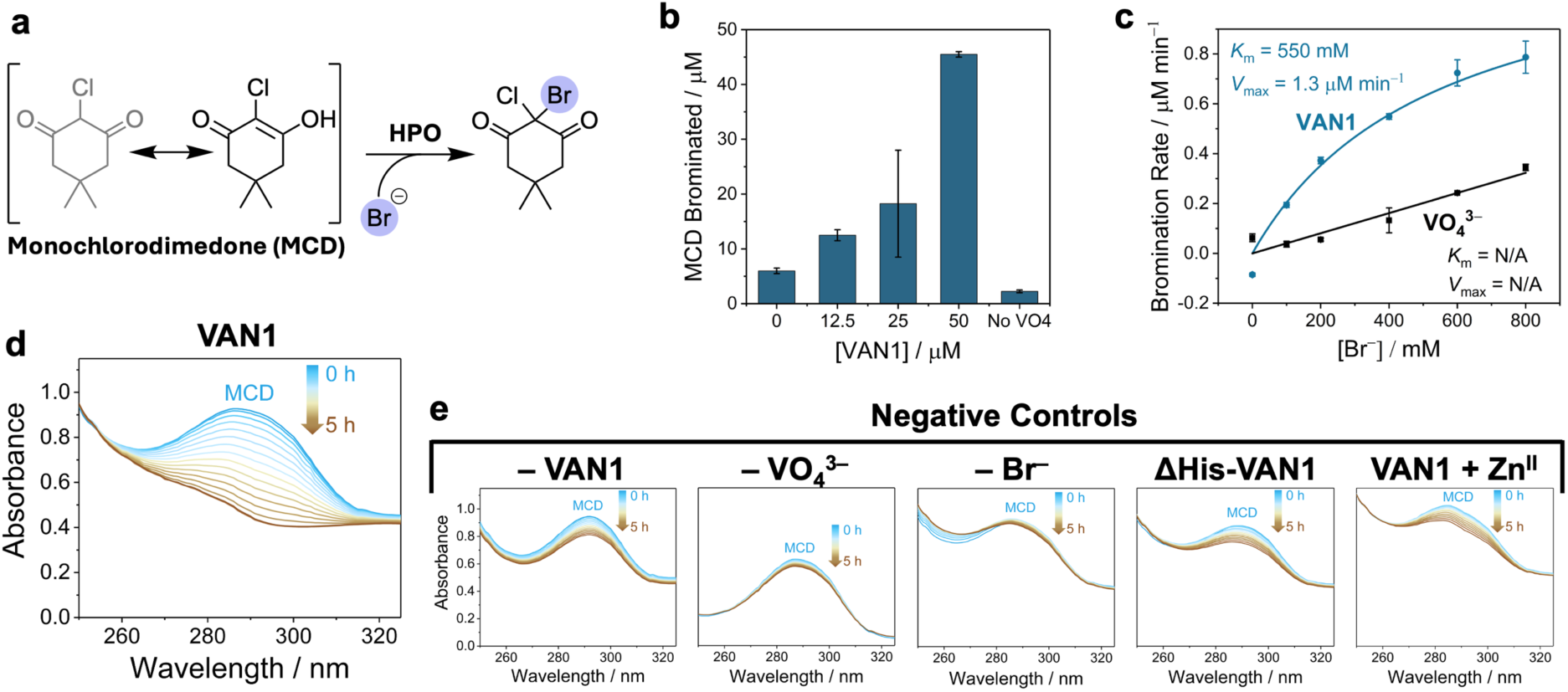
Bromoperoxidase activity of VAN1. (a) Monochlorodimedone (MCD) reaction scheme. (b) Batch MCD reactions (50 µM MCD) at increasing concentrations of VAN1 (reactions carried out in duplicate). See Fig. S3a–e for absorbance traces. (c) MCD assays at increasing [Br^−^] show that VAN1 follows Michaelis-Menten kinetics while control assays without VAN1 vary linearly with [Br^−^]. (d) VAN1 timecourse assay showing the complete bromination of 100 µM MCD over 4.7 h (see Fig. S3f–i for additional kinetic data). (e) Negative control MCD assays run in parallel with panel d without added VAN1, VO_4_^3−^, or Br^−^, using the negative-control VAN1 variant, ΔHis-VAN1, which should not bind VO_4_^3−^, or with 1 mM ZnSO_4_ which acts as a competitive inhibitor by preventing VO_4_^3−^ binding at the active site.

We carried out extensive control experiments in parallel with the VAN1 assays to test whether the activity observed was due to VAN1 catalysis occurring at the designed VO_4_^3−^ binding site (Fig. 3e, Figs. S3–4). Experiments without VAN1 displayed only a low background bromination rate catalyzed by VO_4_^3−^ in solution, while assays lacking VO_4_^3−^ or Br^−^ resulted in essentially no change in the concentration of MCD (Fig. 3e, Fig. S3d–e). Having confirmed that VAN1, VO_4_^3−^ and Br^−^ are each required for activity, we designed a negative-control VAN1 variant to verify whether the designed VO_4_^3−^ binding site was responsible for the bromination. This variant, ΔHis-VAN1, had the two putative VO_4_^3−^ binding His residues (H93/H151) replaced with Ala, resulting in a protein that should not bind VO_4_^3−^. When assayed alongside VAN1, which completely brominated the MCD substrate after 4.7 h (Fig. 3d), the ΔHis-VAN1 variant did not show activity above the background level (Fig. 3e), strongly suggesting that the designed VO_4_^3−^ binding site is the VAN1 active site. To independently confirm this, we performed experiments using Zn^II^ as a competitive inhibitor. AlphaFold3^36^ (AF3) predicted that VAN1 binds Zn^II^ at the designed VO_4_^3−^ binding site using H93 and H151 along with two nearby Glu residues (E89/E149) (Fig. S4a). Competitive metal binding titrations confirmed that VAN1 tightly binds one equivalent of Zn^II^ with an average *K*_d_ = 10 nM (Fig. S4b–c), and a Zn-VAN1 crystal structure (see below) confirmed the Zn^II^ binding location. In the presence of Zn^II^, VAN1 no longer catalyzed the bromination of MCD (Fig. 3e, Fig. S4d), confirming that the designed VO_4_^3−^ binding pocket is the VAN1 active site. Control experiments showed that high concentrations of Zn^II^ did not affect the background bromination rate catalyzed by VO_4_^3−^ in solution (Fig. S4d).

While VAN1 was much more active than all controls over a period of several hours, we consistently observed a lag phase (Fig. S5a) which we hypothesized was due to oxidation of the protein methionine residues by the oxidized bromide species, forming methionine sulfoxide (MetO) and temporarily decreasing the reaction rate by competing with the MCD substrate (Fig. S5b). To test whether Met oxidation occurs, we ran MCD assays for 5 hours before isolating the proteins and performing mass spectrometry (MS) analysis. MS analysis confirmed a 64–66 Da mass increase, consistent with the oxidation of all four Met residues (Fig. S5c). To test if Met oxidation affected the bromination assays, we carried out a control experiment in which VAN1 was replaced with a 4-fold molar excess of free Met, which did not significantly affect the background reaction rate (Fig. S4d). Furthermore, we do not expect that Met oxidation negatively affects VAN1 activity or folding as the residues are located far from the active site.

### Solution Characterization of VAN1

Having established that VAN1 displays bromoperoxidase activity, we performed extensive physical characterization of the protein. MS and analytical ultracentrifugation (AUC) confirmed that VAN1 and ΔHis-VAN1 have the expected size (20.1 kDa) and exist as monomers in solution (Fig. S6a–c). Denaturation experiments revealed that VAN1 is highly thermally and chemically stable, maintaining 60–70% of its folded structure at 98 °C, only completely denaturing in the presence of 6 M guanidine, and fully refolding upon cooling or removal of guanidine (Fig. S6d–g). Furthermore, VAN1 maintained its folded structure when dissolved in mixed aqueous/organic solutions ranging from 10–100% (v/v) of methanol, ethanol, or acetonitrile (Fig. S6h–k). The ΔHis-VAN1 variant had a virtually identical circular dichroism (CD) spectrum and exhibited similar thermostability compared to VAN1 (see below), indicating that the negative-control variant was a faithful representation of the VAN1 scaffold in solution. Thus, the lack of bromination activity for ΔHis-VAN1 is solely due to the mutations at the active site.

### Structural Characterization of Metal-bound VAN1

Following extensive screening, apo-VAN1 crystallized after several weeks in the presence of 1.2–1.6 M succinate (pH = 7.0–8.5). Dozens of datasets were collected, however, solving the structure proved challenging because all VAN1 crystals exhibited pseudo-merohedral twinning,^37, 38^ mimicking a higher-order symmetry space group. To overcome this, we soaked apo-VAN1 crystals with 2 mM ZnCl_2_ and solved the Zn-VAN1 structure using a combination of molecular replacement and single-wavelength anomalous diffraction (MR-SAD). The apo-VAN1 AF3 prediction was used as the search model, and the anomalous Zn signal provided independent validation of the protein placement and improved the phasing. Six independent Zn-VAN1 datasets were merged to yield a final structure at 2.69 Å resolution (Fig. 4a–d) containing eight copies of Zn-VAN1 in the asymmetric unit (ASU).

**Fig. 4.**
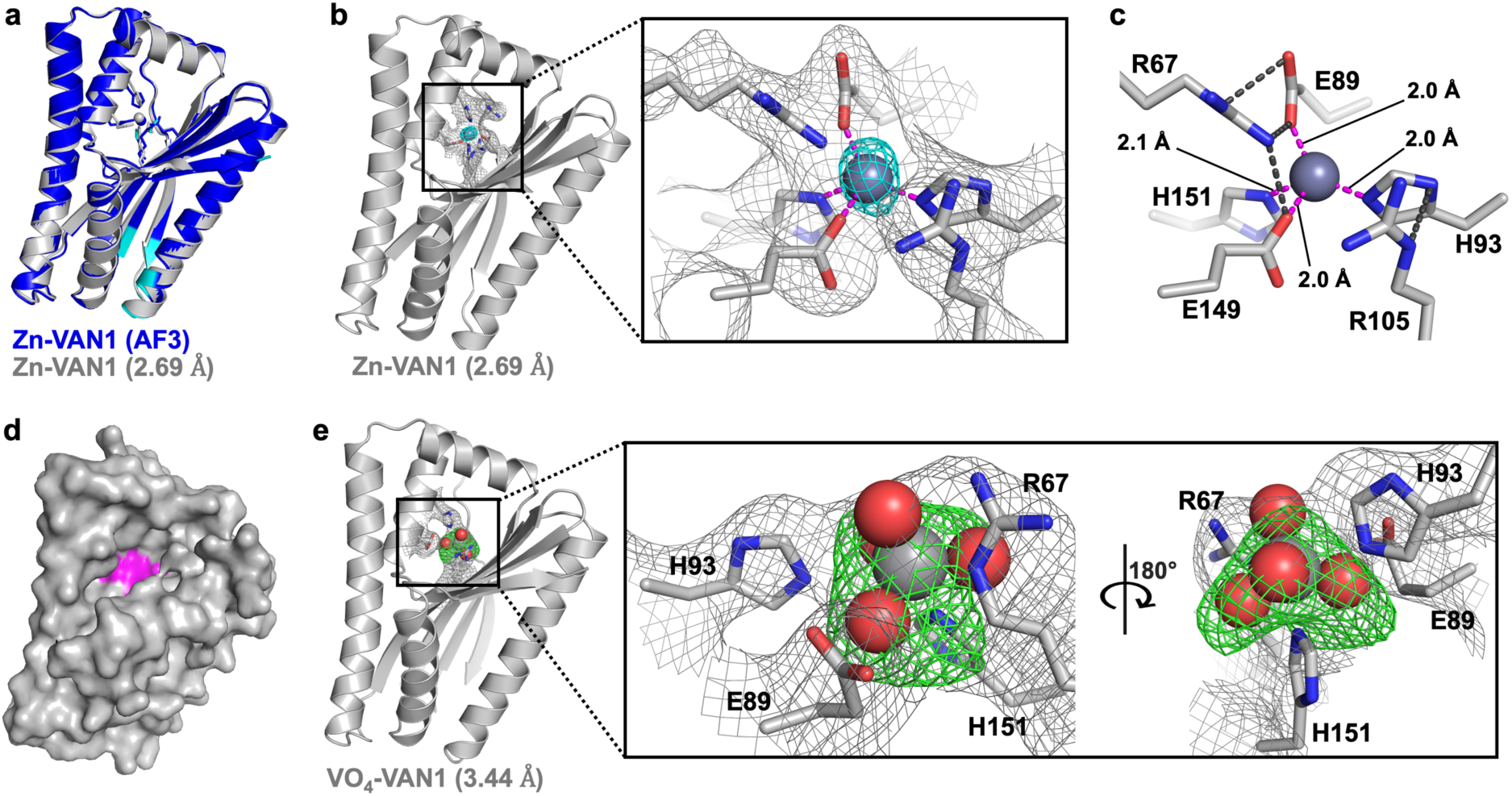
Structural characterization of Zn-VAN1 and VO_4_-VAN1. (a) AlphaFold3 prediction of Zn-VAN1 superposed with the X-ray crystal structure of Zn-VAN1 determined at 2.69 Å. The RMSD values range from 0.6–0.8 Å due to slight differences between protein chains in the asymmetric unit. (b) Crystal structure of Zn-VAN1 with the Zn^II^ binding site enlarged. The 2*F*_o_ – *F*_c_ map is shown at 1.0σ (grey mesh) and the anomalous Zn map is shown at 8σ (cyan mesh). (c) Stick view of the Zn^II^ binding site showing that Zn^II^ is bound in a distorted tetrahedral geometry with R67 providing hydrogen bonding interactions to help stabilize E89 and E149 and R105 H-bonding to H93. (d) Surface view of VAN1 with the designed VO_4_^3–^ binding site/Zn^II^ binding site colored magenta to show that the active site is partially buried within the protein but that it remains accessible to solvent and small molecule substrates. (e) Crystal structure of VO_4_-VAN1 (3.44 Å) showing VO_4_^3–^ bound at the designed binding site (2*F*_o_ – *F*_c_ = 1.0σ). To verify the presence of VO_4_^3–^, we generated a difference map (*F*_o_ – *F*_c_; 3.0σ) with VO_4_^3–^ removed from the model (green mesh): 5 out of 8 chains in the ASU possessed strong electron density at the designed VO ^3–^ binding site in the calculated difference map with VO_4_^3–^ omitted.

The Zn-VAN1 crystal structure was highly similar to the AF3 prediction (RMSD = 0.6– 0.8 Å; Fig. 4a). The Zn primary coordination sphere consisted of the same four residues predicted by AF3, including the two conserved His residues for VO_4_^3−^ binding, H93 and H151, and two nearby Glu residues (E89 and E149). A notable difference from the AF3 prediction was the orientation of a secondary-sphere arginine residue (R67). In the crystal structure, this arginine adopts a conformation which provides hydrogen bonding interactions that stabilize the Zn-coordinating E89 and E149 residues (Fig. 4b,c). Surface representations of the structure highlight a distinct, solvent-exposed pocket leading to the metal center (Fig. 4d), confirming that the designed active site is contained within a protected cavity while remaining sufficiently exposed for small-molecule substrate access. The most prominent protein packing interaction formed between two helices of adjacent VAN1 molecules in which the interface was composed entirely of Ala residues (Fig. S7). This alanine zipper motif drew our attention to the overall hydrophobicity of VAN1. Notably, hydrophobic residues were not limited to the protein core but also populated the protein exterior (Fig. S7), resulting in an overall hydrophobic protein with a grand average of hydropathicity (GRAVY) score of –0.19 and an aliphatic index of 96.6. This is significantly more hydrophobic than the natural chloroperoxidase from which we derived the starting motif (GRAVY = –0.441; aliphatic index = 72.1).

Obtaining a high-resolution structure for VO_4_-VAN1 was more challenging. After numerous attempts, the structure was solved at 3.44 Å by soaking apo-VAN1 crystals with 17 mM Na_3_VO_4_ (Fig. 4e). Excitingly, electron density consistent with VO_4_^3−^ occupancy was observed at the designed VO_4_^3−^ binding site. Since V does not produce a strong anomalous signal, we generated an omit difference map (with VO_4_^3−^ excluded from the model) to verify the presence of VO_4_^3−^, which showed that five out of eight protein chains in the asymmetric unit possessed strong density at the designed binding site (Fig. 4e). Due to the low resolution of the data, we could not determine the precise primary coordination sphere of the metal cofactor. However, our findings are consistent with VO_4_^3−^ binding at the designed binding site and supported by the inactive ΔHis-VAN1 variant and the inhibitory effect of Zn^II^ (Fig. 3e), which binds at the same site (Fig. 4a–c).

### Rational Mutagenesis for Improved VO_4_^3−^ Binding and Catalysis

Although VAN1 possesses halogenation activity, unsurprisingly, the bromination rates are low relative to natural haloperoxidases. Furthermore, relatively high concentrations of VO_4_^3−^ (> 2 mM) are required to consistently achieve bromination rates significantly above the background, suggesting weak binding affinity for VO_4_^3−^. With the goal of improving both VO_4_^3−^ binding affinity and bromination activity, we introduced rational mutations at the VAN1 active site.

In natural V haloperoxidases, a single His residue directly coordinates the V atom and the VO_4_^3−^ oxygens are stabilized by H-bonding interactions within an overall positively charged active site.^19^ Upon inspecting the Zn-VAN1 and VO_4_-VAN1 crystal structures (Fig. 4) and the VAN1 AF3 prediction with docked VO_4_^3−^ (Fig. 5a), we postulated that the two Zn-coordinating Glu residues (E89/E149) may impede VO_4_^3−^ binding through both steric and repulsive ionic effects. To test this hypothesis, we designed four single-mutant variants of VAN1 in which the two Glu residues were mutated to either Ala, to ameliorate steric clashes with VO_4_^3−^, or to Asn, which is structurally closer to Glu while being one carbon atom shorter and neutrally charged. All four variants were expressed, and VAN1^E89N^, VAN1^E149A^, and VAN1^E149N^ were purified in high yield. VAN1^E89A^ was much less soluble, and only a small amount of the protein was obtained after two expression attempts. We characterized each variant using CD spectroscopy to confirm folding and test thermostability (Fig. 5b). Both E89 variants (VAN1^E89A^ and VAN1^E89N^) exhibited high thermostability similar to VAN1 and ΔHis-VAN1, remaining ∼70% folded at 98 °C (Fig. 5b). In contrast, VAN1^E149A^ and VAN1^E149N^ were less thermostable, remaining only ∼50 % folded at 98 °C (Fig. 5b), suggesting that Glu149 provides an important role in stabilizing the VAN1 tertiary structure.

**Fig. 5.**
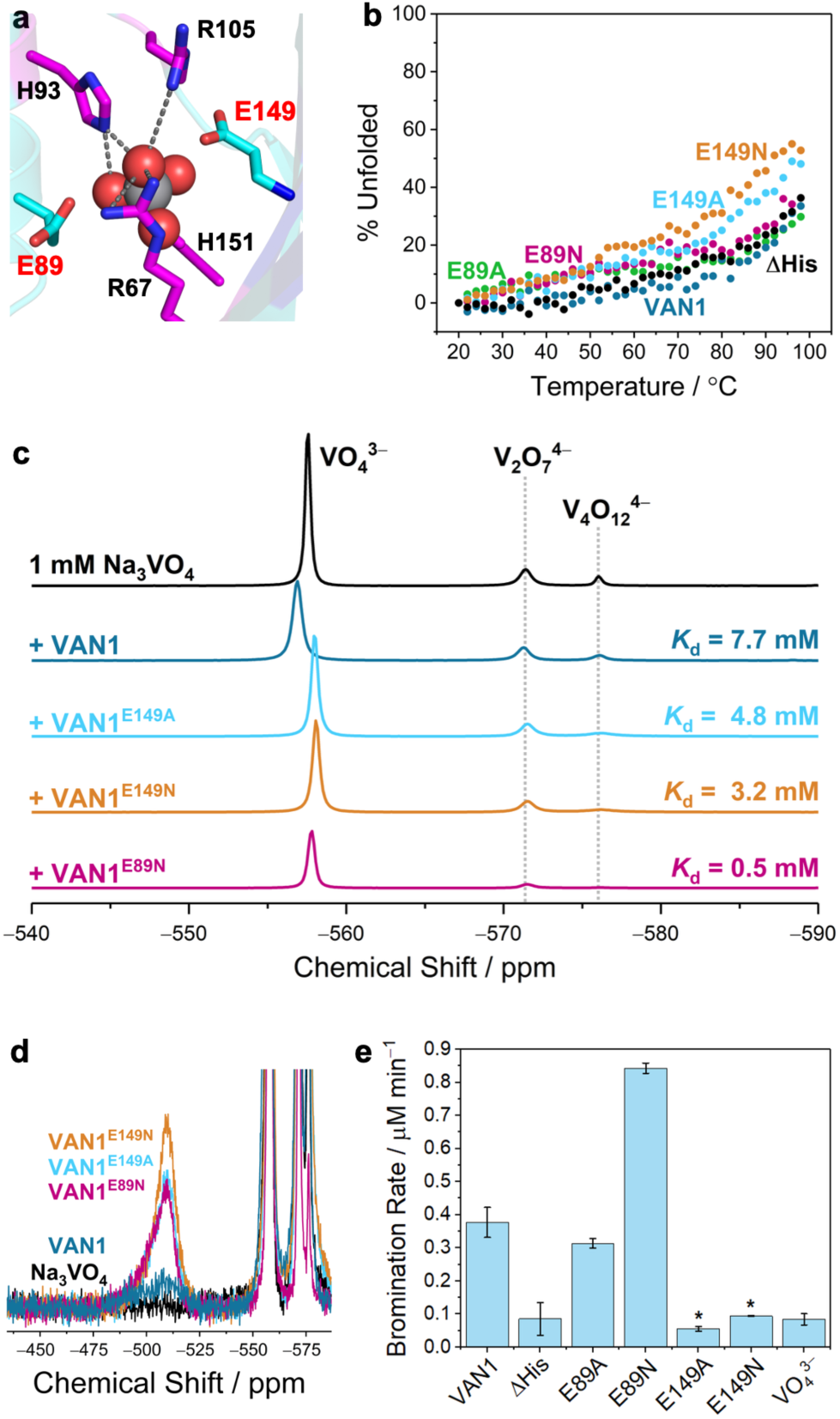
Rational mutagenesis of VAN1 and quantification of VO_4_^3–^ binding affinities. (a) AF3 prediction of VAN1 with VO_4_^3–^ docked into the active site. The two mutated Glu residues (E89/E149) predicted to decrease VO_4_^3–^ affinity are colored cyan. (b) Thermal denaturation curves for VAN1, the VAN1 variants, and ΔHis-VAN1. (c) ^51^V NMR quantification of residual solvated ^51^V species (pH = 7.4) with and without added protein to quantify VO_4_^3–^ binding affinities. Repeat ^51^V NMR experiments and concentrations used to calculate *K*_d_ values are given in Table S1. (d) Zoomed-in view showing the peaks assigned to the protein-VO_4_^3–^ complexes at 1 mM Na_3_VO_4_ and ∼1 mM of each protein. (e) Bromination rates obtained from MCD assays (100 µM MCD) for each of the VAN1 variants.

To quantify the VO_4_^3−^ affinity of VAN1 and its variants, we first employed inductively coupled plasma mass spectrometry (ICP-MS). ICP-MS analysis confirmed that VAN1 binds Zn^II^ ions while the negative control ΔHis-VAN1 variant does not (Fig. S8a). However, required washing steps and low concentrations of protein and VO_4_^3−^ precluded VO_4_^3−^ quantification. Thus, we turned to ^51^V NMR, a well-established method for studying protein-VO_4_^3−^ interactions.^22, 39, 40, 41^ A 1 mM solution of Na_3_VO_4_ yielded the expected peaks at pH 7.4, with monomeric VO_4_^3−^ at –558 ppm and minor peaks for pyrovanadate (V_2_O_7_^4−^) and tetravanadate (V_4_O_12_^4−^) at –571 and –576 ppm, respectively (Fig. 5c). VAN1 and its variants were incubated with Na_3_VO_4_ for 1 h, and the decrease in the solvated ^51^V signal was used to calculate binding affinities (Fig. 5c, Table S1). As expected, VAN1 exhibited a low affinity for VO_4_^3−^ (*K*_d_ = 7.7 mM).

In contrast, all three tested variants displayed improved VO_4_^3−^ affinities (Fig. 5c, Table S1), confirming that both Glu residues (E89/E149) negatively impact VO_4_^3−^ binding. The two E149 variants (VAN1^E149A^ and VAN1^E149N^) showed moderately improved VO_4_^3−^ affinities (*K*_d_ = 4.8 and 3.2 mM, respectively), whereas replacing the Glu89 residue with Asn (VAN1^E89N^) resulted in a 59% reduction in the free VO_4_^3−^ signal (at 1 mM protein) corresponding to a *K*_d_ = 0.5 mM, a 15-fold improvement over VAN1 (Fig. 5c, Table S1). Importantly, these measurements were made on proteins with identical mass (20.1 kDa) at the same concentrations (∼1 mM). Thus, the sixfold range in the magnitude of the ^51^V signal decrease (9%, 11%, 16% and 59% for VAN1, VAN1^E149A^, VAN1^E149N^ and VAN1^E89N^, respectively; Table S1) can be attributed to the active-site substitutions alone. Alongside the decrease in the free ^51^V signal, we observed a broad ^51^V-protein signal at –509 ppm (Fig. 5d), consistent with previously reported ^51^V-protein species.^33, 34^ ^35^ Due to the broad nature of this peak, coupled with complex quadrupolar ^51^V relaxation effects,^40^ the protein-VO_4_^3−^ signal was used solely for qualitative confirmation of VAN1-VO_4_^3−^ complex formation.

Having confirmed that the rational mutations improved VO_4_^3−^ binding affinity, we next tested whether this translated to improved haloperoxidase activity. Consistent with their lower thermostability, both VAN1^E149A^ and VAN1^E149N^ were less stable under assay conditions (Fig. 5e). VAN1^E89A^ was also less stable than VAN1 or VAN1^E89N^ and did not exhibit clear improvements in activity (Fig. 5e). Notably, VAN1^E89N^, the variant with 15-fold higher VO_4_^3−^ binding affinity, catalyzed the bromination of MCD at a 2.6-fold greater rate than VAN1 (Fig. 5e). Given the spread in the measured *K*_d_ values (Table S1), VAN1 is expected to be 32–36% metalated under the assay conditions (4 mM VO_4_^3−^) compared to 82–92% for VAN1^E89N^, corresponding to a predicted rate enhancement of 2.3–2.9-fold. The measured 2.6-fold rate increase falls at the midpoint of this range, indicating that the improvement in activity is quantitatively accounted for by the increase in VO_4_^3−^ occupancy at the designed site. These results demonstrate that the VAN1 scaffold is amenable to engineering, making it a valuable new starting point for V-HPO mechanistic studies.

### Bromination of Tyrosine, Catalytic Temperature Dependence, and Substrate Selectivity

To further investigate the catalytic properties of VAN1 and VAN1^E89N^, we measured their activity with more challenging substrates than MCD. We pursued two alternative substrates with activated phenolic rings, 4-hydroxybenzyl alcohol (4-HBA) and *L*-tyrosine. While the enzymes brominated both 4-HBA and tyrosine, additional products generated from 4-HBA complicated quantification. Consequently, we focused on *L*-tyrosine as a representative aromatic substrate, which can undergo sequential bromination to yield 3-bromotyrosine and 3,5-dibromotyrosine (Fig. 6a).^42^ We quantified the brominated products using reversed-phase HPLC (Fig. 6b).

**Fig. 6.**
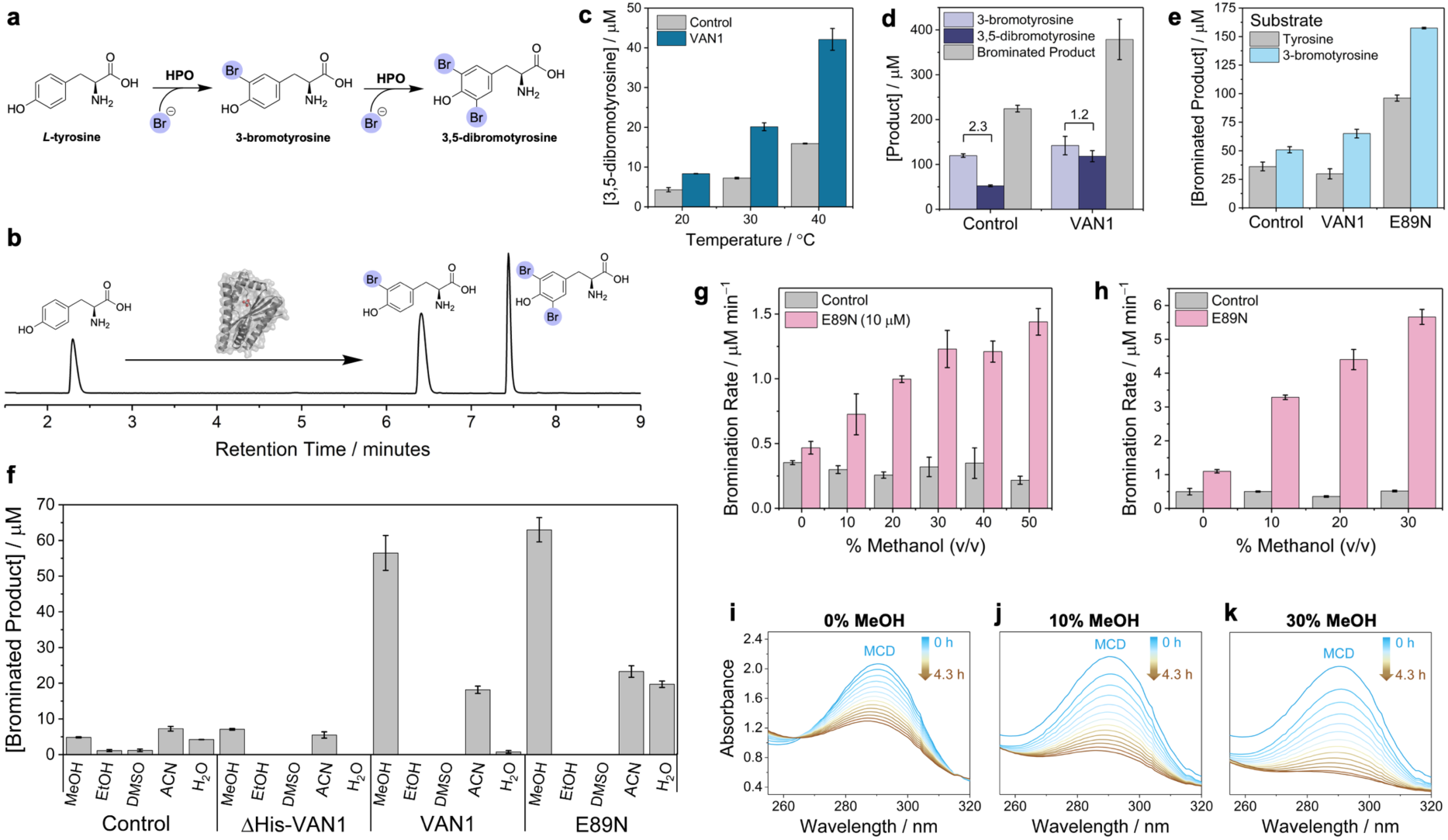
Investigating the temperature dependence, substrate selectivity, and solvent tolerance of VAN1 and VAN1^E89N^. (a) Reaction scheme showing the bromination of tyrosine to yield 3-bromotyrosine and 3,5-dibromotyrosine. (b) HPLC quantification of tyrosine, 3-bromotyrosine, and 3,5-dibromotyrosine. (c) Concentrations of 3,5-dibromotyrosine after 6 h under standard aqueous assay conditions at different reaction temperatures with 1 mM *L*-tyrosine as the substrate. (d) Reaction products after 21 h for the 20 °C assay in panel c showing that VAN1 produces more 3,5-dibromotyrosine than expected suggesting VAN1 selectivity for the 3-bromotyrosine substrate. (e) Aqueous assays comparing the total brominated product after 6 h using 1 mM tyrosine or 1 mM 3-bromotyrosine as the substrate. Where tyrosine was the substrate, [brominated product] = [3-bromotyrosine] + 2[3,5-dibromotyrosine]; for reactions with 3-bromotyrosine, [brominated product] = [3,5-dibromotyrosine]. (f) Tyrosine bromination assays in which the reactions were diluted 2-fold in either methanol, ethanol, DMSO, acetonitrile, or water to measure enzyme activity in 50% of each solvent. The reaction conditions were identical to those in panels b–e prior to dilution in each solvent. (g–h) MCD bromination rates (400 µM MCD) for 10 µM (g) or 50 µM (h) VAN1^E89N^ in the presence of 0–50% methanol. (i–k) MCD kinetic scans showing the decrease in the MCD absorbance signal (290 nm) over time for VAN1^E89N^ at 0–30% methanol. Note: reactions labeled “control” are identical to other conditions but without added protein.

Given the high thermostability of VAN1, we carried out assays at five temperatures up to 80 °C. However, under the highly oxidizing assay conditions, temperatures exceeding 40 °C resulted in the formation of oligomerized tyrosine products that precluded quantification. Therefore, triplicate reactions at 20, 30, and 40 °C were quenched after 6 h and the products quantified by HPLC. As anticipated, bromination activity increased significantly as the temperature increased (Fig. 6c, Fig. S9a–b). However, the distribution of brominated products differed significantly between the VAN1 assays and the VO_4_^3−^ controls. VAN1 yielded only slightly more 3-bromotyrosine but produced ∼2-fold more 3,5-dibromotyrosine relative to VO_4_^3−^ controls (Fig. S9a–b), suggesting that VAN1 may selectively brominate 3-bromotyrosine over tyrosine. This trend was also observed after 21 h at 20 °C, yielding a 3-bromotyrosine/3,5-dibromotyrosine ratio of 2.3 for the VO_4_^3−^ control versus 1.2 for VAN1 (Fig. 6d). To directly test substrate selectivity, we performed assays using 1 mM tyrosine or 1 mM 3-bromotyrosine and analyzed the total brominated product after 6 h (Fig. 6e). In contrast to VO_4_^3−^ controls, which consistently produced similar amounts of product for both substrates, VAN1 and VAN1^E89N^ each displayed significantly higher activity with 3-bromotyrosine than with tyrosine (Fig 6e, Fig. S9c). To quantify selectivity while accounting for inherent reactivity differences between substrates, we calculated the ratio of the enzyme-catalyzed product yields for each substrate after normalizing each value to the VO_4_^3−^ control yields (roughly proportional to the ratio of apparent rate enhancements for each substrate assuming rate linearity over 6 h).^43^ Over two independent experiments, VAN1 displayed an average 1.8-fold (± 0.2) preference for 3-bromotyrosine over tyrosine. VAN1^E89N^ had a slightly lower ratio in favor of 3-bromotyrosine (1.2-fold), however this may reflect the higher overall activity of VAN1^E89N^ and the decreased reaction rate after 6 h. One possible explanation for this selectivity is that the more hydrophobic 3-bromotyrosine has greater affinity for the hydrophobic interior of VAN1.

### VAN1 and VAN1^E89N^ Display Enhanced Bromination Activity in Organic Solvents

Because VAN1 is a relatively hydrophobic protein (Fig. S7) that remains folded in several water-miscible organic solvents (Fig. S6h–k), we assessed the bromination activity of VAN1 and VAN1^E89N^ in 50% (v/v) of methanol, ethanol, dimethyl sulfoxide (DMSO), and acetonitrile. Assay solutions were diluted 2-fold into each solvent alongside a water-diluted control, quenched after 6 h, and analyzed by HPLC (Fig. 6f). The VAN1 and VAN1^E89N^ reactions diluted with methanol generated significantly more product than any other condition, including water-diluted controls, indicating that 50% methanol enhances the bromination activity of both enzymes. Control assays without added protein or with ΔHis-VAN1 each produced very little product, confirming that the VAN1 VO_4_^3−^ active site is responsible for the observed catalysis in 50% organic solvent. The addition of acetonitrile also significantly enhanced the activity of VAN1, whereas 50% acetonitrile with VAN1^E89N^ yielded similar activity to the water-diluted assays. Despite the higher activities observed in methanol and acetonitrile, no bromination was observed in assays diluted with ethanol or DMSO. Interestingly, VAN1 and VAN1^E89N^ displayed similar activity in methanol and acetonitrile, even though VAN1^E89N^ is far more active when diluted with water (Fig. 6e) or under standard aqueous conditions (Fig. 5e). These observations may result from the mild enzyme aggregation observed following the addition of organic solvents, or from altered VAN1-VO_4_^3−^ binding under these conditions.

To confirm the solvent effects, we repeated assays with methanol, ethanol, and water, and included 3-bromotyrosine as an alternative substrate (Fig. S9d). Consistent with our previous results, methanol-diluted assays yielded dramatically higher activity, while the addition of ethanol resulted in no brominated product. Compared to aqueous conditions, VAN1 displayed lower selectivity for 3-bromotyrosine in methanol, consistent with the hypothesis that enzyme selectivity may be driven by increased affinity for the more hydrophobic 3-bromotyrosine. The lack of activity in DMSO can be attributed to DMSO oxidation by the brominating species, in the form of either HOBr or an active-site-bound Br^+^ species, to form dimethyl sulfone,^44^ effectively quenching the reaction. Similarly, ethanol reacts with Br^+^ species or HOBr to form ethyl hypobromite, which decomposes to acetaldehyde and Br^−^.^45, 46, 47^ In contrast, methyl hypobromite is more kinetically stable than ethyl hypobromite^48, 49^ due to the higher p*K*a of the methanol *a*C, which must be deprotonated to drive elimination of Br^−^. Thus, if methyl hypobromite forms, it may serve as an oxidized bromine carrier,^48^ unlike ethanol, which neutralizes enzyme-generated HOBr. Acetonitrile likely does not react with HOBr, and therefore the elevated activity observed in both acetonitrile and methanol suggests that activity enhancement in mixed aqueous/organic solvent is a general feature of VAN1. Possible explanations include the enzyme adopting a more favorable conformation, increased protein flexibility to accommodate substrate, improved affinity for VO_4_^3−^, or suppression of background HOBr quenching.

To better assess the solvent enhancement effects without diluting all reaction components two-fold, we carried out MCD kinetic assays under standard conditions in the presence of 0–50% methanol (Fig. 6g–k). Preliminary experiments revealed that VAN1^E89N^ slowly aggregated at concentrations ≥40% methanol, but reducing the enzyme concentration to 10 µM mitigated this effect. At 10 µM, the aqueous VAN1^E89N^ control displayed only marginally higher activity compared to the background, whereas bromination rates increased substantially in the presence of methanol until around 30% at which point the activity increase plateaued. Standard VAN1^E89N^ assays with 30% methanol catalyzed bromination at >10-fold higher rates than the background and ∼8-fold higher than the equivalent aqueous condition, resulting in a background-corrected enzyme turnover frequency of 0.11 min^−1^ (Fig. 6h) and completely brominating 400 µM MCD in 4.3 h.

## Conclusions

The existence of only two known V-dependent enzymes, despite high marine abundance of VO_4_^3−^ and the catalytic versatility of synthetic V complexes, exemplifies the limited reaction space explored by natural evolution. This limitation may result from the inherent difficulty of binding bulky, anionic oxometalates that rely primarily on weaker hydrogen-bonding and electrostatic interactions rather than the strong coordination bonds utilized by cationic transition metals. VAN1 reflects these principles through its low binding affinity for VO_4_^3−^ (*K*_d_ = 7.7 mM) in stark contrast with the unintended tight binding of Znᴵᴵ (*K*_d_ = 10 nM), which binds the designed active site by recruiting two nearby Glu residues alongside the two conserved His residues. This 10^6^-fold affinity difference demonstrates the relative ease with which binding sites can be designed (or form spontaneously) for biologically prevalent transition metals (e.g., Fe, Cu, Zn) compared to oxometalates (Fig. 1a) and may help explain why catalytically versatile and abundant metals like V and Mo are underutilized in biology.

In addition to binding VO_4_^3−^, VAN1 represents the first *de novo* protein with halogenase activity. Maintaining only four conserved active-site residues at the VO_4_^3−^ binding site, the activity of VAN1 confirms that the highly conserved structural features of natural V-HPOs are not strictly necessary to achieve basal catalytic activity. However, the low activities of both VAN1 and VO_4_^3−^-reconstituted bacterial acid phosphatases^20^ relative to natural V-HPOs reflect the importance of evolved functional optimization and a precisely arranged secondary coordination sphere. Despite the initial low affinity of VAN1 for VO_4_^3−^, rational mutagenesis yielded the VAN1^E89N^ variant, which has 15-fold higher affinity (*K*_d_ = 0.5 mM) and a 2.6-fold higher catalytic rate, demonstrating the ease with which the VAN1 scaffold can be tuned. Finally, the high stability and significantly enhanced bromination activity of VAN1 in mixed aqueous-organic solvents represent significant advantages over most natural enzymes for industrial applications. With the rapid advancement of ML tools for protein design and structure prediction, the vast reaction space currently unoccupied by natural metalloenzymes presents a massive opportunity. Designed proteins can now serve as tunable ligand platforms for metal-catalyzed, new-to-nature chemistry. Ultimately, this work provides a new starting point for both mechanistic studies and engineering of V-HPOs and expands the scope of *de novo* enzyme design to include challenging C–X bond formation.

## Author Contributions

R.A.H. conceived the project and carried out the protein design and biochemical/structural characterization. R.A.H. and L.A.C. expressed/purified protein and performed CD spectroscopy, competitive binding titrations, and crystallized VAN1. R.A.H. and K.J.A. performed ^51^V NMR experiments under the supervision of K.J.G. R.A.H. and F.A.T. directed the project and wrote the manuscript. All authors approved the final version of the manuscript.

## Acknowledgements

This work was supported by the National Institutes of Health (Grant R01-GM138884 and R35-GM163543 to F.A.T.). R.A.H. was supported by an NIH Ruth L. Kirschstein Postdoctoral Fellowship (F32-GM157932). X-ray diffraction experiments were performed at the Advanced Light Source (5.0.1 and 8.3.1 beamlines) supported by the DOE-BES (DE-AC02-05CH11231) and NIH (P30-GM124169-01), and the Stanford Synchrotron Radiation Lightsource supported by DOE-BES (DE-AC02-76SF00515).

## Data availability

Atomic coordinates and structure factors for Zn-VAN1 and VO_4_-VAN1 have been deposited in the Protein Data Bank under accession codes [39IG] and [39JA], respectively. All other data supporting the findings of this study are available within the paper and its Supporting Information, or from the corresponding authors upon reasonable request.

## Code availability

The RFdiffusion and ProteinMPNN scripts used to generate the VAN1 backbone and sequences are provided as Supporting Data 1 and 2. RFdiffusion, ProteinMPNN, AlphaFold2 and AlphaFold3 are publicly available from their respective developers.

## Methods

### Plasmid Construction

Proteins were expressed with a C-terminal 6x-His-tag with a Tobacco Etch Virus (TEV) protease cleavage site between the 6x-His-tag and protein (Table S2). In all cases, the 6x-His tag was cleaved before characterization to prevent non-specific metal binding. Genes were obtained from Integrated DNA Technologies and assembled into a pET20b(+) plasmid via Gibson assembly. The Gibson assembly product was transformed into *E. coli* XL1-Blue cells, plated onto LB-agar plates containing 100 mg L^−1^ ampicillin, and grown overnight at 37 °C. Single colonies were used to inoculate 5 mL of LB media with 100 mg L^−1^ ampicillin and grown overnight. The next day, plasmid DNA was extracted using a Miniprep kit (NEB) and sequenced using Sanger sequencing by Azenta Life Sciences.

### Protein Expression

Plasmids containing the gene to be expressed were transformed into *E. coli* BL21(DE3) cells and plated onto LB/Agar plates containing 100 mg L^−1^ ampicillin and grown overnight at 37 °C. Single colonies were used to inoculate starter cultures containing 5 mL LB media with 100 mg L^−1^ ampicillin and grown overnight at 37 °C. 2.8 L flasks containing 1 L of autoinduction TBM-5052 media^50^ and 100 mg L^−1^ ampicillin were inoculated with 1 mL of the starter culture, shaken at 200 rpm, and grown for 16–18 h at 37 °C. Cells were harvested by centrifugation at 5000 rpm (10 min) and stored overnight at –80 °C.

Frozen cell pellets were thawed and resuspended in 200 mL of 20 mM MOPS (pH = 8.0) buffer with 300 mM NaCl and lysed by sonication in the presence of a small amount of lysozyme and DNAse. Throughout purification, PO_4_^3−^ buffer was avoided to prevent binding at the designed VO_4_^3−^ binding site. Cell lysate was centrifuged at 18,000 rpm for 25 min and loaded onto a gravity flow column with Ni-NTA resin which had been pre-equilibrated with the MOPS buffer. The protein-loaded column was washed twice with 200 mL of buffer then successive 200 mL fractions containing increasing concentrations of imidazole (30 mM, 40 mM, 50 mM, 50 mM, 150 mM, 500 mM). The proteins generally eluted in the 50 mM and 150 mM imidazole fractions. All fractions were collected separately and analyzed by SDS-PAGE. Fractions containing pure protein were concentrated and buffer exchanged to remove excess imidazole using a 10 kDa Amicon membrane. 1 mM ethylenediaminetetraacetic acid (EDTA) was added when the imidazole concentration was below 10 mM, and the 6x-His tag was cleaved from the protein by addition of 1.3–2.6 mg of TEV protease and overnight incubation at 4 °C. Following cleavage, the protein was buffer exchanged to remove the EDTA (< 1 µM final concentration). The protein was concentrated to 100 mL and passed over the Ni-NTA column. The flow through and two washes were collected to obtain the pure cleaved protein while the TEV protease remained bound to the Ni-NTA as it possessed a 6x-His tag. The fractions (including 150 and 500 mM imidazole fractions) were collected and analyzed by SDS-PAGE. Fractions containing pure, cleaved protein were incubated for >1 h with 1 mM EDTA, concentrated to 1–2 mL, and buffer exchanged into 20 mM MOPS (pH 7.4) buffer to remove residual EDTA (final [EDTA] < 1 µM). Protein concentration was determined using the average value of two independent Bradford assays. Protein stocks were divided into single-use aliquots, flash frozen, and stored at −80 °C.

### Haloperoxidase Assays

#### Monochlorodimedone (MCD) Assays

MCD assays were carried out in either 1.5 mL Eppendorf tubes (for batch reactions) or 96-well plates (for kinetic assays). Standard assay conditions were as follows: 50–400 µM MCD (indicated in figure captions), 4 mM Na_3_VO_4_, 400 mM NaBr, 4 mM H_2_O_2_, 53 mM MES buffer solution (pH = 5.2). All values indicate the final assay concentrations. Batch reactions were incubated at room temperature (21 °C) and the assays were diluted and analyzed using an Agilent Cary 60 UV-Vis spectrometer in a quartz cuvette with a pathlength of 1 cm. Kinetic assays were carried out at 25 °C using a Tecan Infinite M Nano^+^ plate reader with Greiner UV-Star microplate flat-bottom 96-well plates. Bromination of MCD product was determined by monitoring the decrease in absorbance at 290 nm (ε_290_ = 20200 M^−1^ cm^−1^).^34^ For Zn^II^ inhibition experiments, a final concentration of 1 mM ZnSO_4_ was added after initiating the reaction. All reactions were carried out in triplicate unless otherwise noted.

#### Tyrosine and 3-bromotyrosine bromination assays

Tyrosine assays were carried out in 200 µL PCR tubes using the same standard conditions as the MCD assays except that 1 mM of the tyrosine or 3-bromotyrosine substrate was used. Reactions (100 µL) were incubated at room temperature unless otherwise noted, in which case they were incubated in a thermocycler at the specified temperature. Reactions in mixed aqueous/organic solvent used the same conditions as aqueous assays except that after initiating the reaction by addition of enzyme, the entire assay solution was diluted 2-fold with the specified solvent. Reactions were quenched by successive additions of 20 mM sodium thiosulfate (10 min incubation) to react with residual HOBr and H_2_O_2_ followed by 10 U of catalase (15 min incubation) to remove any remaining H_2_O_2_. To precipitate soluble VAN1 enzyme, 100 mM ZnSO_4_ (final concentration) was added^51^ and the solution was incubated for 30 min. Samples were centrifuged for 5 min at 8000 rpm and the supernatant was analyzed by HPLC for brominated products. All reactions were carried out in triplicate.

### Competitive Metal Binding Titrations

Competitive metal-binding titrations were performed using 5 μM protein mixed with 5 μM of the metal-sensitive dye indicator, Mag-Fura-2 in a chelated buffer containing 20 mM MOPS and 150 mM NaCl (pH 7.4). Titrations were carried out in 3 mL of solution in a quartz cuvette with a path length of 1 cm. UV-Vis spectra were recorded using an Agilent Cary 60 UV-Vis spectrometer, and 3–6 μL aliquots of 1 mM ZnCl_2_ were added. The solution was allowed to equilibrate with stirring for 10 minutes after each ZnCl_2_ addition, after which the UV-Vis spectrum was measured. The increase in concentration of metal-bound Mag-Fura-2 was monitored by plotting the A_337 nm_/A_365 nm_ ratio and fitting the curve using Dynafit^52^ as described previously^53^ using a dissociation constant of 3.70 x 10^-8^ M for Mag-Fura-2.^54^

### Analytical Ultracentrifugation (AUC)

Sedimentation velocity measurements were carried out using a Beckman XL-A ultracentrifuge using an An-60 Ti rotor. The VAN1 concentration was 110 μL dissolved in 20 mM MOPS (pH 7.4) chelated buffer. Scans (500–1000) were obtained at 25 °C at 41,000 rpm. The data were analyzed using SEDFIT^55^ software using the following parameters: buffer density = [1.0022] g/mL; buffer viscosity = [0.00902] poise; partial specific volume = [0.74844].

### Mass Spectrometry

Electrospray ionization mass spectrometry (ESI-MS) as performed using an LTQ XL linear ion trap mass spectrometer (Thermo Fisher Scientific) with a heated electrospray ionization (HESI) source. Proteins were diluted to 50 μM in HPLC grade water containing 0.2% formic acid; samples were then diluted 2-fold in acetonitrile (final sample contained 25 μM protein with 0.1% formic acid and 50% acetonitrile). The capillary temperature was maintained at 275 °C, samples were injected with a flow rate of 5 µL min^−1^ and auxiliary gas flow rate of 12 arb. The spray voltage was 5 kV, and the source temperature was 100 °C. For each sample, 200 spectra were collected and averaged using the Thermo Xcalibur software. Averaged spectra were deconvoluted using UniDec.^56^

### Circular Dichroism (CD) Spectroscopy

CD spectra were acquired on an Applied Photophysics Chirascan CD spectrometer at a protein concentration of 10 μM in a chloride-free buffer solution of 20 mM borate (pH 7.4) in a 2 mm pathlength quartz cuvette. CD spectra were collected with 0.5 nm intervals and an averaging time of 1 s at 25 °C. Thermal denaturation studies were performed by monitoring the average CD signal from 220–224 nm with 2 °C intervals and a 120 s temperature equilibration time between steps. GuHCl denaturation studies were performed by monitoring the CD signal at 220–224 nm between 0 and 8 M GuHCl.

### Inductively Coupled Plasma Mass Spectrometry (ICP-MS)

Proteins were diluted to a concentration of 10 μM and incubated with 10 μM of either ZnCl_2_ or Na_3_VO_4_ (500 μL total volume) for 1 h. Protein samples were then exhaustively buffer exchanged (>15000-fold dilution) using Amicon centrifugal spin columns (10 kDa cutoff) into a 20 mM HEPES buffer solution (pH = 7.5). Proteins were concentrated to a volume of 100 μL, and 50 μL of 30% (v/v) H_2_O_2_ and 50 μL of 1% HCl were added to each sample. The acidified protein solutions were incubated at 60 °C for 1 h using a thermal cycler. Following incubation, the samples were diluted with a 1% HCl solution to a total volume of 5 mL. Samples were analyzed by ICP-MS using a PerkinElmer NexION 2000 instrument at the California NanoSystems Institute at the University of California, Los Angeles.

### Structure Determination by X-Ray Diffraction

VAN1 was crystallized by sitting-drop vapor diffusion at room temperature. The sitting drop was made by adding 1 μL of precipitant solution to 1 μL of protein. The reservoir contained 200 μL of precipitant solution. VAN1 crystals appeared within several weeks. Metal solutions (either ZnCl_2_ or Na_3_VO_4_) were mixed 1:1 with the precipitant solution, and 1 μL of this solution was mixed into the solution containing VAN1 crystals. The crystals were incubated with metals at room temperature for 2 h. Crystallization conditions are shown in Table S3. Picked crystals were soaked in perfluoro polyether (Hampton) for cryoprotection prior to flash freezing. The X-ray diffraction datasets were obtained at 100 K at the ALS beamlines 5.0.1 and 8.3.1. Datasets were processed using XDS^57^ software. Using the apo-VAN1 AlphaFold prediction as the search model and the Zn^II^ anomalous signal, a combined molecular replacement and single-wavelength anomalous diffraction (MR-SAD) approach was used to solve the structure of VAN1 using Phenix.^58^ Data resolution was truncated based on CC_1/2_ significance (CC_1/2_ > 0.1) and paired-refinement analysis, confirming that inclusion of the high-resolution shell improved overall R_free_ and map quality.^59^ Twinned refinement was carried out using a combination of automated refinement with phenix.refine and manual refinement using Coot.^60^ No bond distance restraints were used for the VO_4_-VAN1 structure. Loose bond distance restraints of 2.05 Å (σ = 0.05 Å) were used for the Zn atoms and coordinating residues in the Zn-VAN1 structure. X-ray data collection and refinement statistics are given in Table S4.

### 51V NMR Spectroscopy

Proteins were mixed with Na_3_VO_4_ (500 μL total volume) in a 20 mM MOPS buffer solution (pH = 7.4), incubated for 1 h, and transferred to Wilmad NMR tubes (7 in; 5 mm OD; 4 mm ID). ^51^V NMR measurements were performed at *B*_0_ = 9.4 T, *ν_L_*(^1^H) = 400.13 MHz using a Bruker Ascend magnet, a Bruker Avance Neo spectrometer, and a 5 mm Bruker BBO probe. 1,843,200 pulse-acquire scans were recorded using a recycle delay of 50 ms (at least 5x the longitudinal relaxation time of the slowest-relaxing species) and a 90-degree pulse of 10.5 µs. The spectrum width was 100 kHz with a pre-scan delay of 5 µs and an acquisition time of 40.96 ms. ^51^V NMR chemical shifts were indirectly referenced to neat VOCl_3_ through the ^13^C resonance of neat tetramethylsilane at −0.71 ppm using a frequency ratio *Ξ* of 26.302948 for ^51^V.^61^ Concentrations of proteins and Na_3_VO_4_ for each measurement are given in Table S1. Dissociation constants were calculated according to: 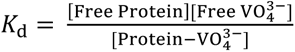. Concentrations of pyrovanadate and tetravanadate are omitted from the equation because they rapidly respond to equilibrium shifts as free VO_4_^3−^ is bound, acting as a VO_4_^3−^ reservoir. Consequently, spectra for proteins with higher VO_4_^3−^ affinities had smaller pyrovanadate and tetravanadate peaks. Only monomeric VO_4_^3−^ was treated as the binding species in this analysis. Binding affinities were determined at pH 7.4, whereas bromoperoxidase assays were carried out at pH 5.2 following a 30 min enzyme-VO_4_^3−^ preincubation at pH 7.4. The occupancy analysis assumes that the relative VO_4_^3−^ affinities of VAN1 and its variants are preserved between these conditions. This assumption is supported by the agreement between the predicted occupancy range and the measured rate enhancement.

### High Performance Liquid Chromatography (HPLC)

Brominated tyrosine products, 3-bromotyrosine and 3,5-dibromotyrosine were quantified using a Thermo Scientific Dionex Ultimate 3000 system (HPLC) equipped with an RS variable wavelength UV-vis detector and an Accucore XL C18 (150 x 4.6 mm) 4 μm particle size column (Catalog No. 74104-154630). Sample injections (10 μL) were performed by an autosampler and run at a flow rate of 1 mL min^−1^ with a mobile phase composed of 99% of buffer A (0.1% formic acid in HPLC-grade water) and 1% of buffer B (0.1% formic acid in acetonitrile). The initial mobile phase (99% A; 1% B) was held for 4 min to elute salts and polar reaction components. From 4–10.5 min, a linear gradient was applied to reach 45% buffer B, during which both 3-bromotyrosine and 3,5-dibromotyrosine eluted. At 10.6 min, buffer B was increased to 100% and held until 17.6 to clean the column. From 17.7–21.7 min, the initial mobile phase (99% A; 1% B) was used to re-equilibrate the column for the next sample. The column oven temperature was 25 °C, and absorbance was measured at 280 nm. Products were quantified using freshly prepared standard curves using analytical standards (MedChemExpress).

## Supporting Figures

**Fig. S1.**
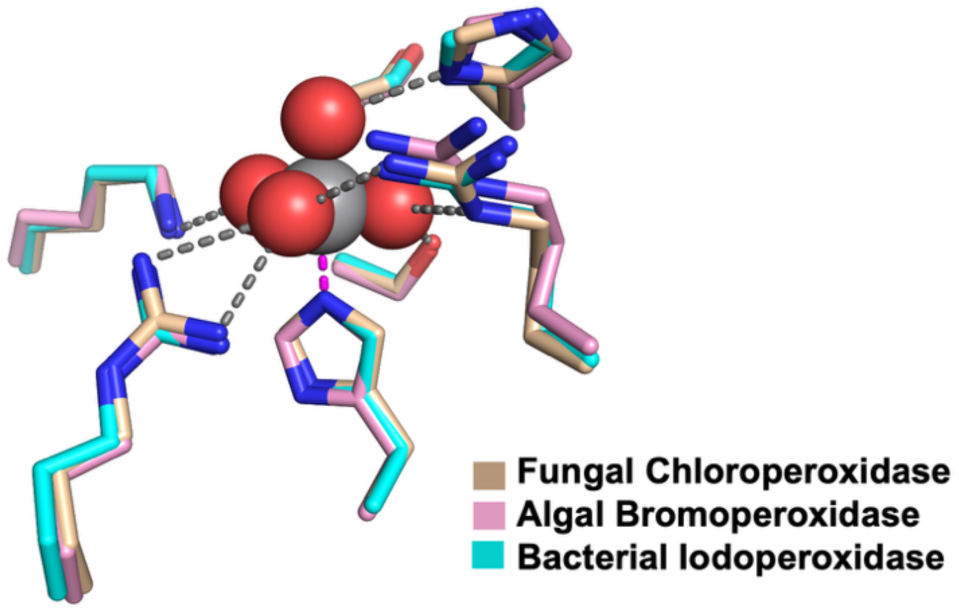
The V-HPO active site is highly conserved as exemplified by the superposition of a fungal chloroperoxidase (wheat; PDB: 1IDQ),^15^ an algal bromoperoxidase (pink; PDB:1QI9),^16^ and a bacterial iodoperoxidase (cyan; PDB: 4CIT).^17^

**Fig. S2.**
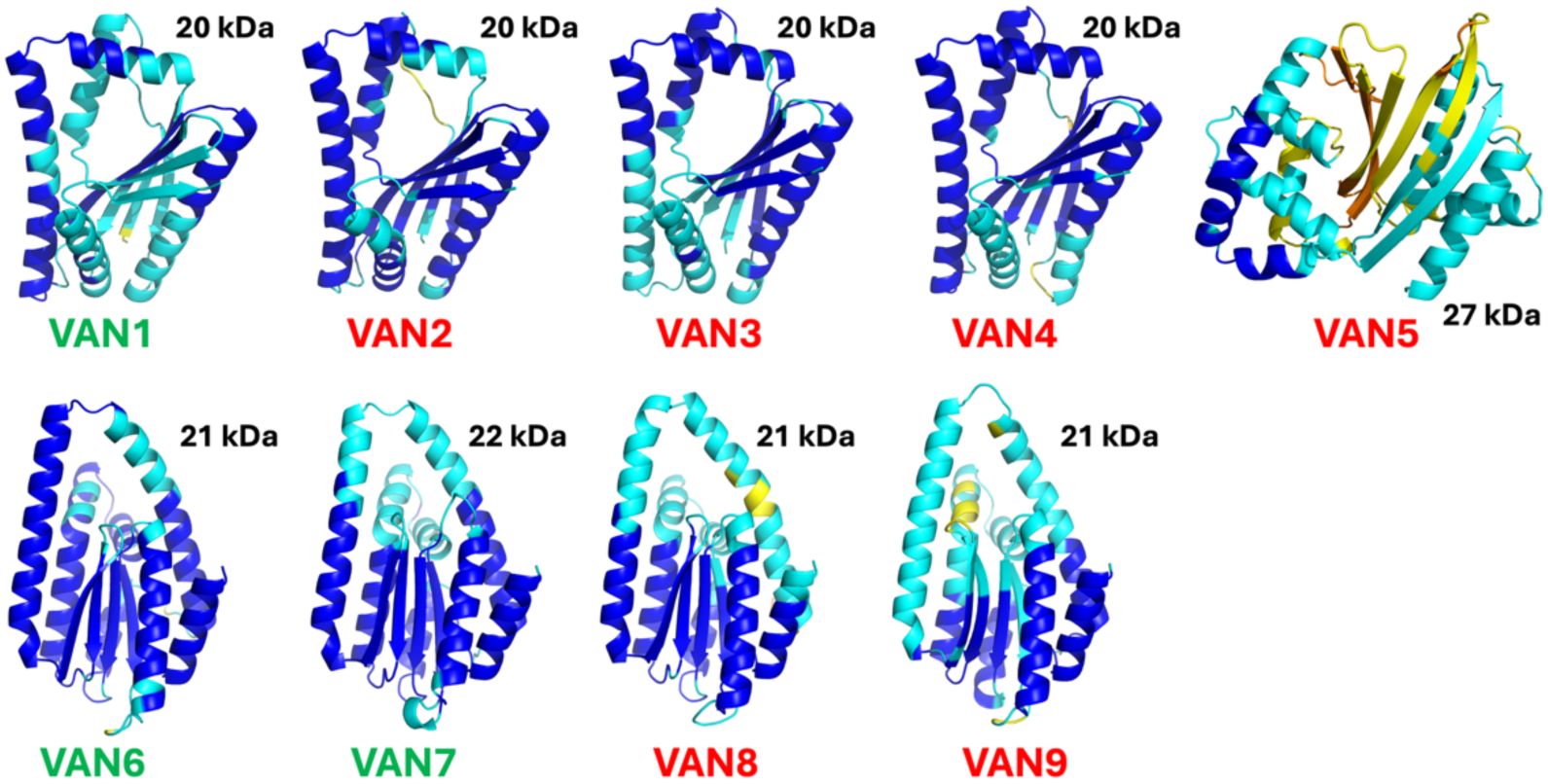
AlphaFold2 predictions of designs which were chosen for expression and characterization. Green-labeled proteins were obtained in high yields, while red-labeled proteins were largely insoluble.

**Fig. S3.**
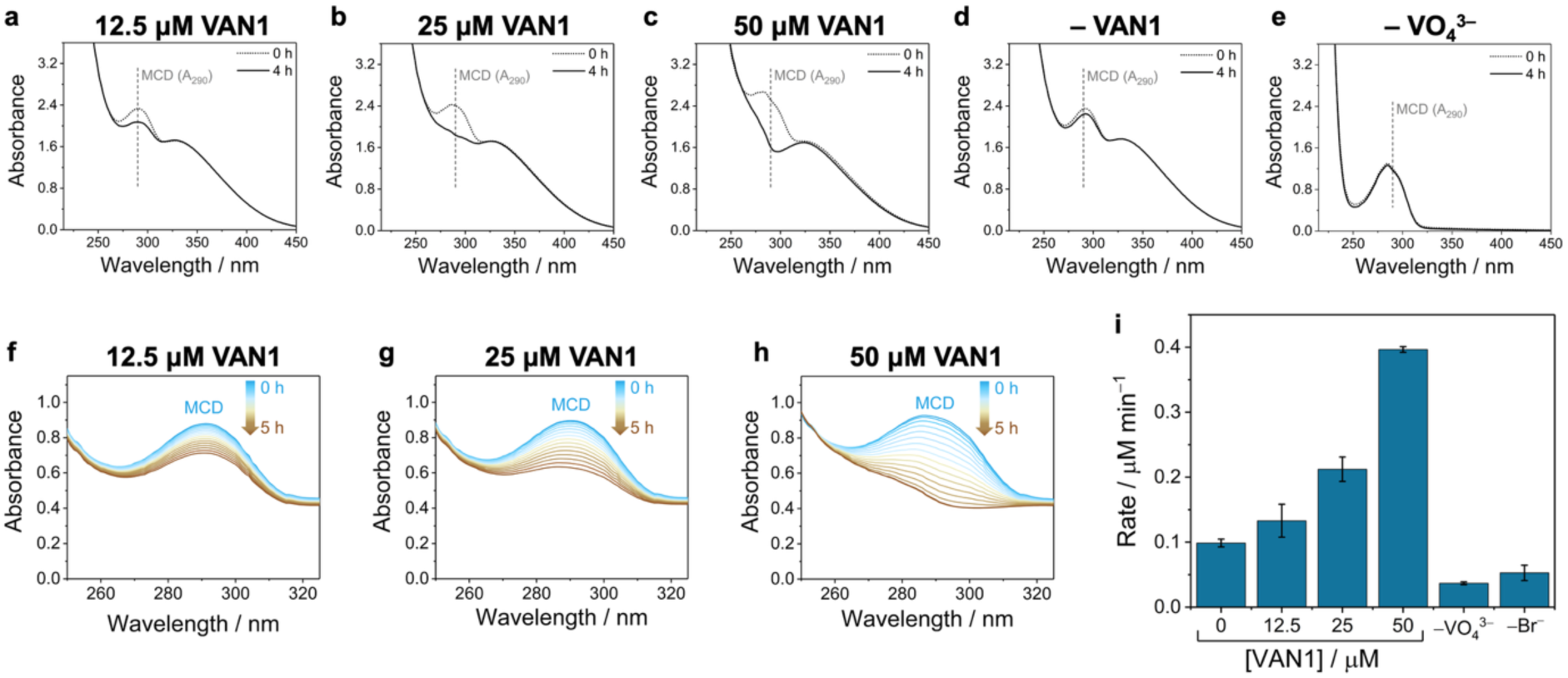
Batch reactions and kinetic MCD assays at different VAN1 concentrations and with additional control experiments. (a–e) Representative UV-vis absorbance traces for the bar chart shown in Fig. 3b (50 μM MCD) with increasing VAN1 concentrations (a–c), without VAN1 added (VO_4_^3–^ only control; d) and without added VO_4_^3–^ (VAN1 only control; e). Samples from each reaction tube were taken at 0 h and 4 h and diluted 5-fold prior to measurement to ensure the absorbance values remained within the linear range of the spectrophotometer. (f–h) Kinetic MCD bromination assays (100 μM MCD) at increasing concentrations of VAN1. Panel h is the same plot shown in Fig. 3d. (i) MCD bromination rates under standard assay conditions (100 μM MCD) at increasing concentrations of VAN1 versus no VO_4_^3–^ and no Br^−^ controls.

**Fig. S4.**
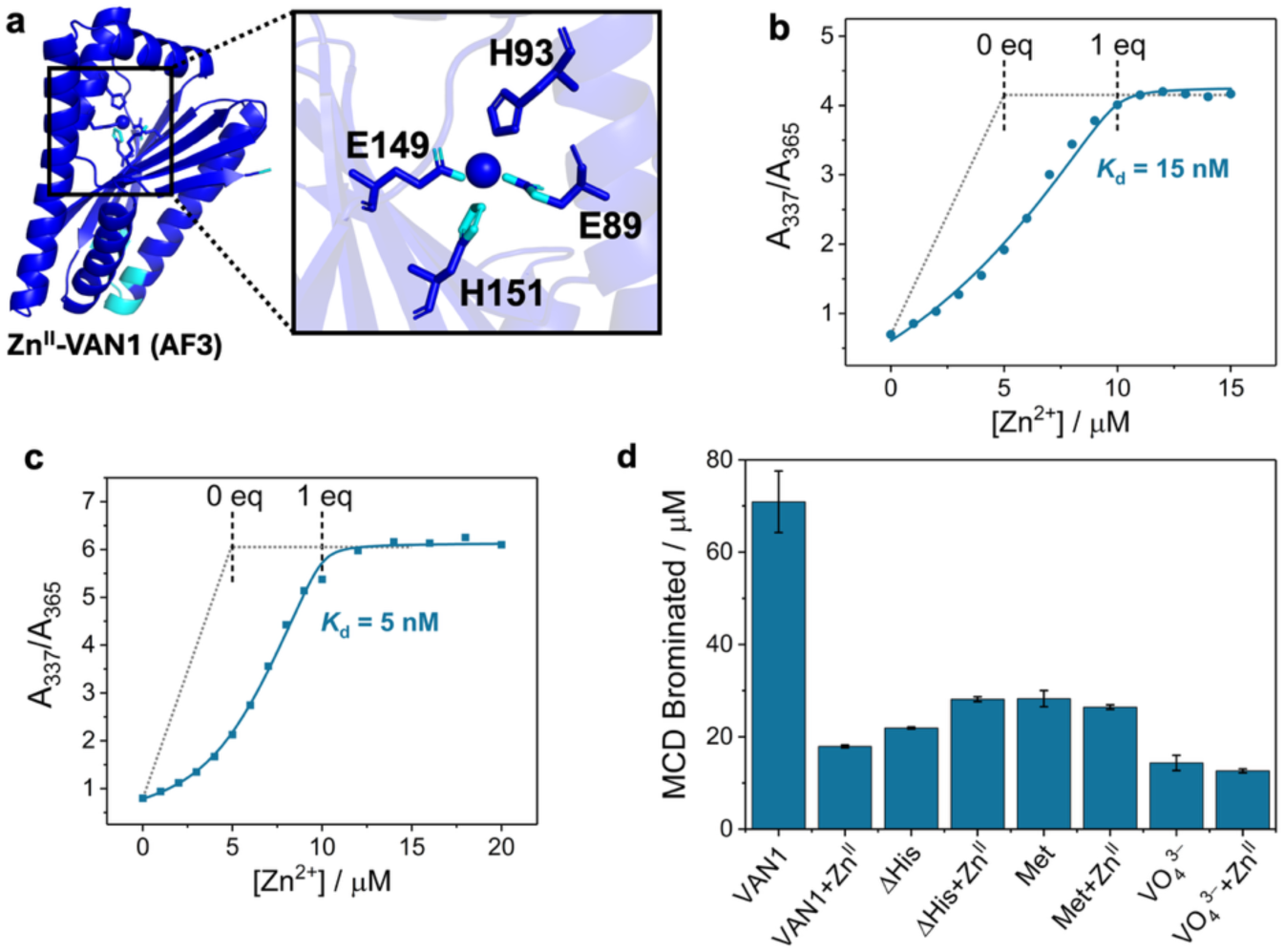
Experiments showing that VAN1 tightly binds one equivalent of Zn^II^ and additional controls with added Zn^II^ and methionine. (a) AF3 predicts that VAN1 binds Zn^II^ at the designed VO_4_^3–^ binding site using the two conserved His residues (H93/H151) and two nearby Glu residues (E89/E149). (b–c) Competitive binding titrations of 5 μM VAN1 against the metal-chelating dye Mag-Fura-2 (5 μM) confirming that VAN1 tightly binds one equivalent of Zn^II^ in solution with an average *K*_d_ = 10 nM. (d) Control experiments testing whether 1 mM added ZnSO_4_ or 200 μM added methionine affect the background bromination rates. In each case, the bromination rate is far lower than VAN1-catalyzed bromination. Assays were carried out for 5 h under standard 96-well conditions (100 μM MCD).

**Fig. S5.**
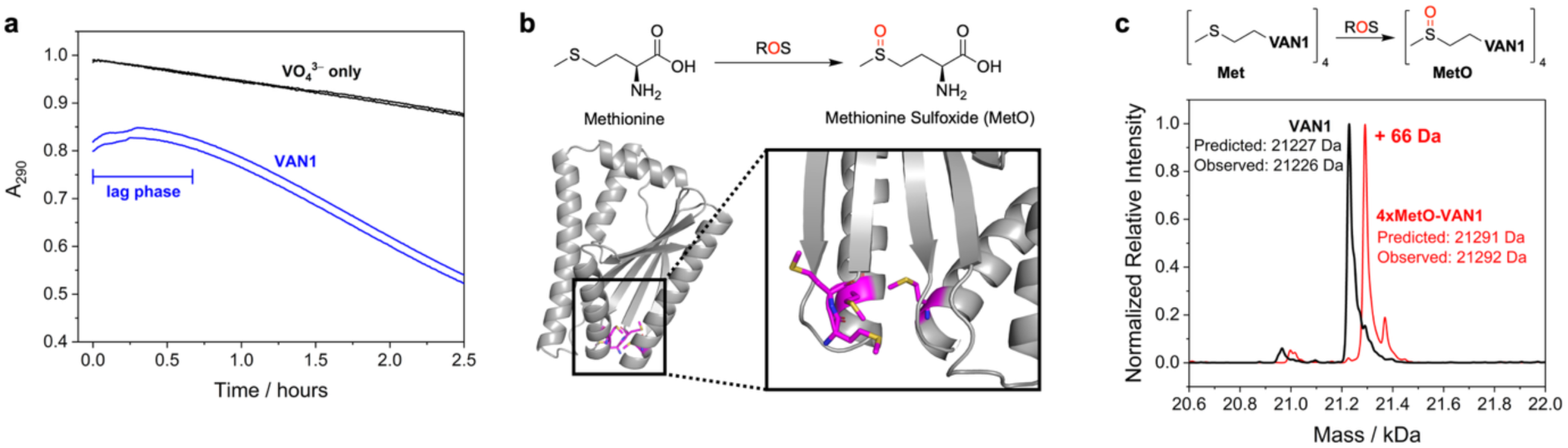
Investigating the VAN1 kinetic lag phase and oxidation of the VAN1 methionine residues under assay conditions. (a) VAN1 exhibits a lag phase during assays that lasts ∼30 min. (b) VAN1 possesses 4 methionine residues, two of which are located on the N-terminus and the other two of which are located near the N-terminus, far from the active site. The reaction scheme shows the oxidation of methionine to methionine sulfoxide (MetO) by reactive oxygen species. (c) Mass spectrometry analysis reveals a +66 Da mass shift for VAN1 which was purified from MCD assays (5 h reaction time) confirming that all four VAN1 Met residues are oxidized to MetO under MCD assay conditions.

**Fig. S6.**
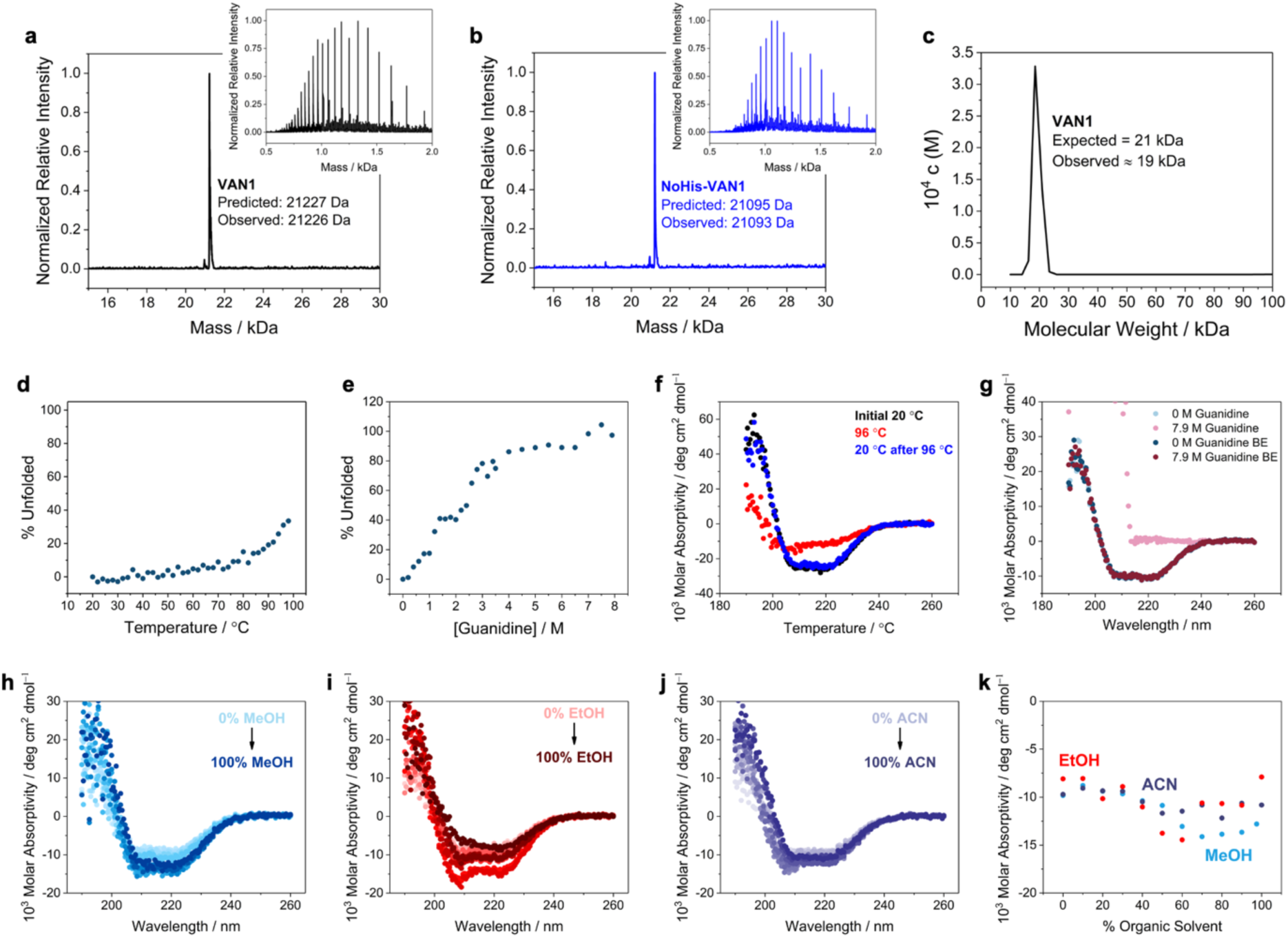
Physical characterization of VAN1. (a–b) Mass spectrometry analysis of VAN1 and ΔHis-VAN1 confirming that they are the expected size. Predicted mass values include the portion of the TEV cleavage site that remained attached to the protein. (c) Sedimentation velocity analysis via analytical ultracentrifugation confirms that VAN1 exists as a monomer in solution. (d) Thermal denaturation of VAN1 (as determined by circular dichroism), showing that VAN1 is highly thermally stable. (e) Chemical denaturation of VAN1 (as determined by circular dichroism), showing that VAN1 is highly chemically stable. (f) Circular dichroism spectra showing that VAN1 can completely refold to its initial tertiary structure after cooling following partial denaturation by heating to 96 °C. (g) Circular dichroism spectra showing that 7.9 M guanidine fully denatures VAN1, but that it refolds into its tertiary structure after the guanidine is removed through extensive buffer exchanging. A control without added guanidine was also buffer exchanged (0 M Guanidine BE) to confirm that the buffer exchanging procedure did not affect VAN1 folding. (h–j) Circular dichroism spectra of VAN1 in the presence of increasing concentrations of different organic solvents (h: methanol; i: ethanol; j: acetonitrile) in 10% increments (from 0% to 100% v/v). VAN1 remains folded in all solvents even at concentrations of 100% solvent. (k) Molar absorptivity measured by the average signal from 220– 224 nm by CD spectroscopy for VAN1 at each concentration of methanol, ethanol, or acetonitrile. Notably, the CD signal becomes more negative for each solvent as the relative fraction increases indicating that the presence of organic solvent may increase the folded state of VAN1 (perhaps by disfavoring looped regions at the termini). Interestingly, this effect decreases for methanol and ethanol at concentrations of solvent >70% by volume.

**Fig. S7.**
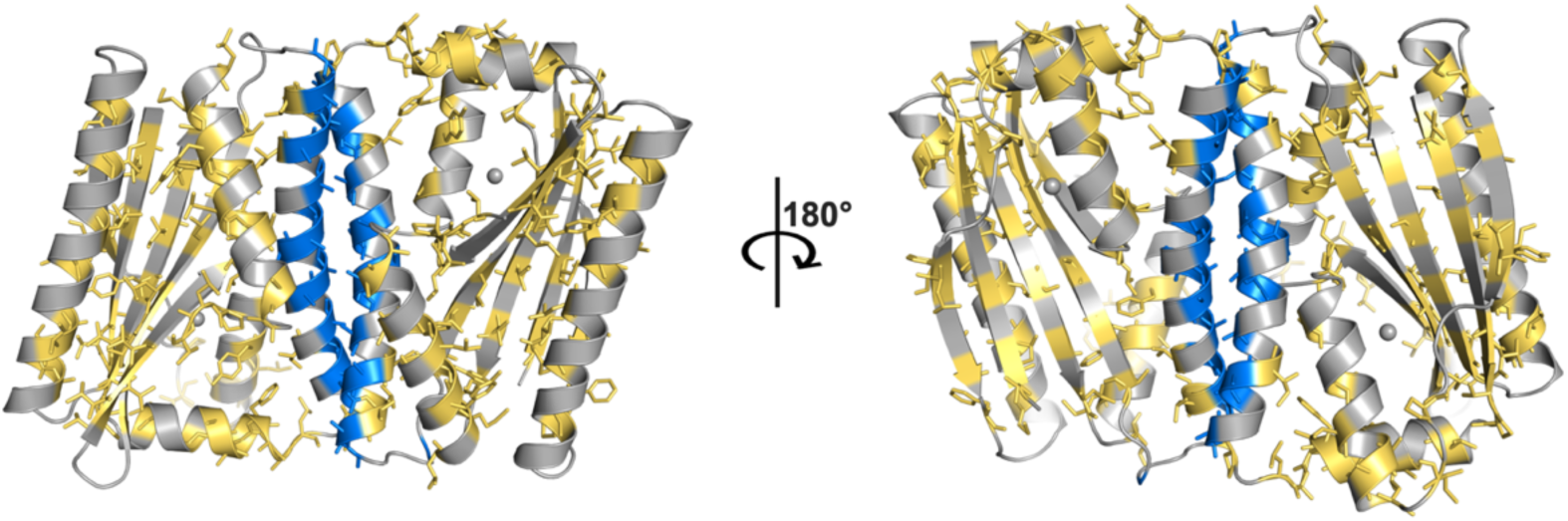
Crystal structure views of a prominent packing interaction in the Zn-VAN1 structure in which two helices interact to form an alanine zipper motif shown in blue. Other hydrophobic residues are shown as yellow sticks, revealing that VAN1 is an overall hydrophobic protein. The presence of many external-facing hydrophobic residues may help to explain why VAN1 is soluble in several organic solvents (Fig. S6h–k) and exhibits enhanced activity in methanol and acetonitrile (Fig. 6f–k).

**Fig. S8.**
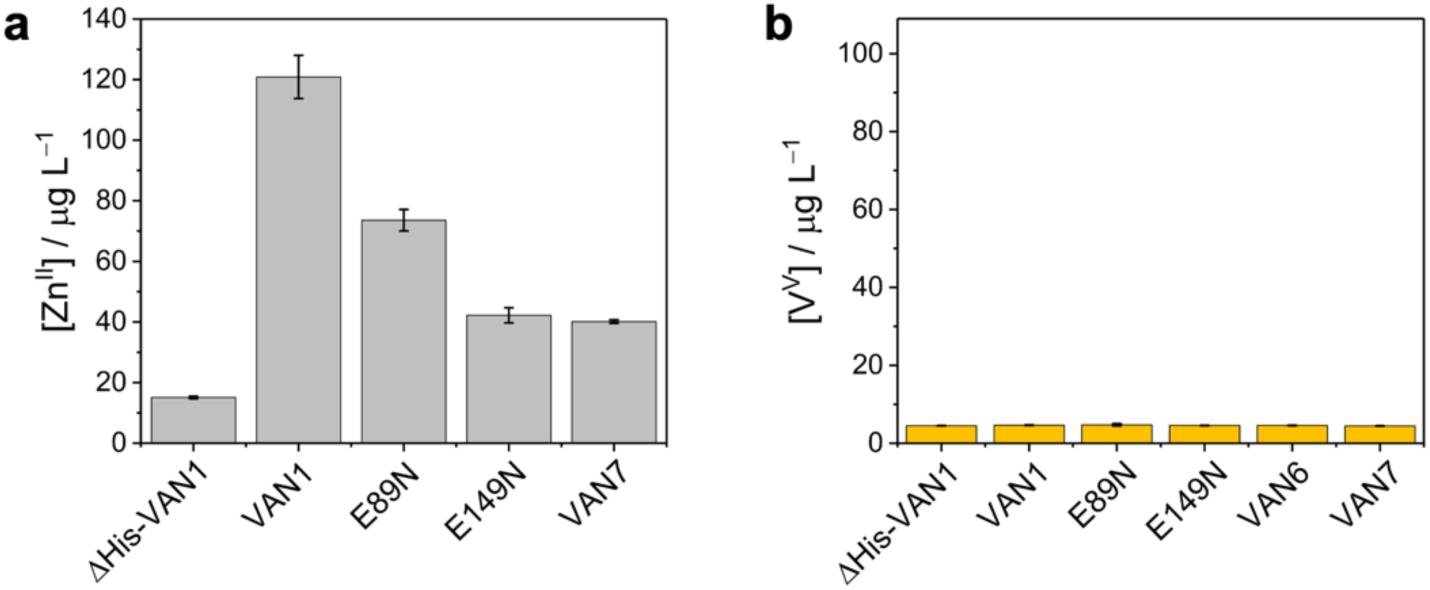
Inductively-coupled plasma mass spectrometry (ICP-MS) analysis. (a) ICP-MS analysis of the presence of Zn for proteins which had been incubated with ZnCl_2_ and extensively buffer exchanged prior to analysis. VAN1 binds Zn while ΔHis-VAN1 does not. VAN1^E89N^, VAN1^E149N^, and VAN7 each bind Zn but with lower affinity than VAN1. (b) ICP-MS analysis measuring the presence of V for proteins incubated with Na_3_VO_4_ extensively buffer exchanged. Due to both the low concentrations of protein and Na_3_VO_4_ and the required washing steps, no VO_4_^3–^ was able to be measured using this method.

**Fig. S9.**
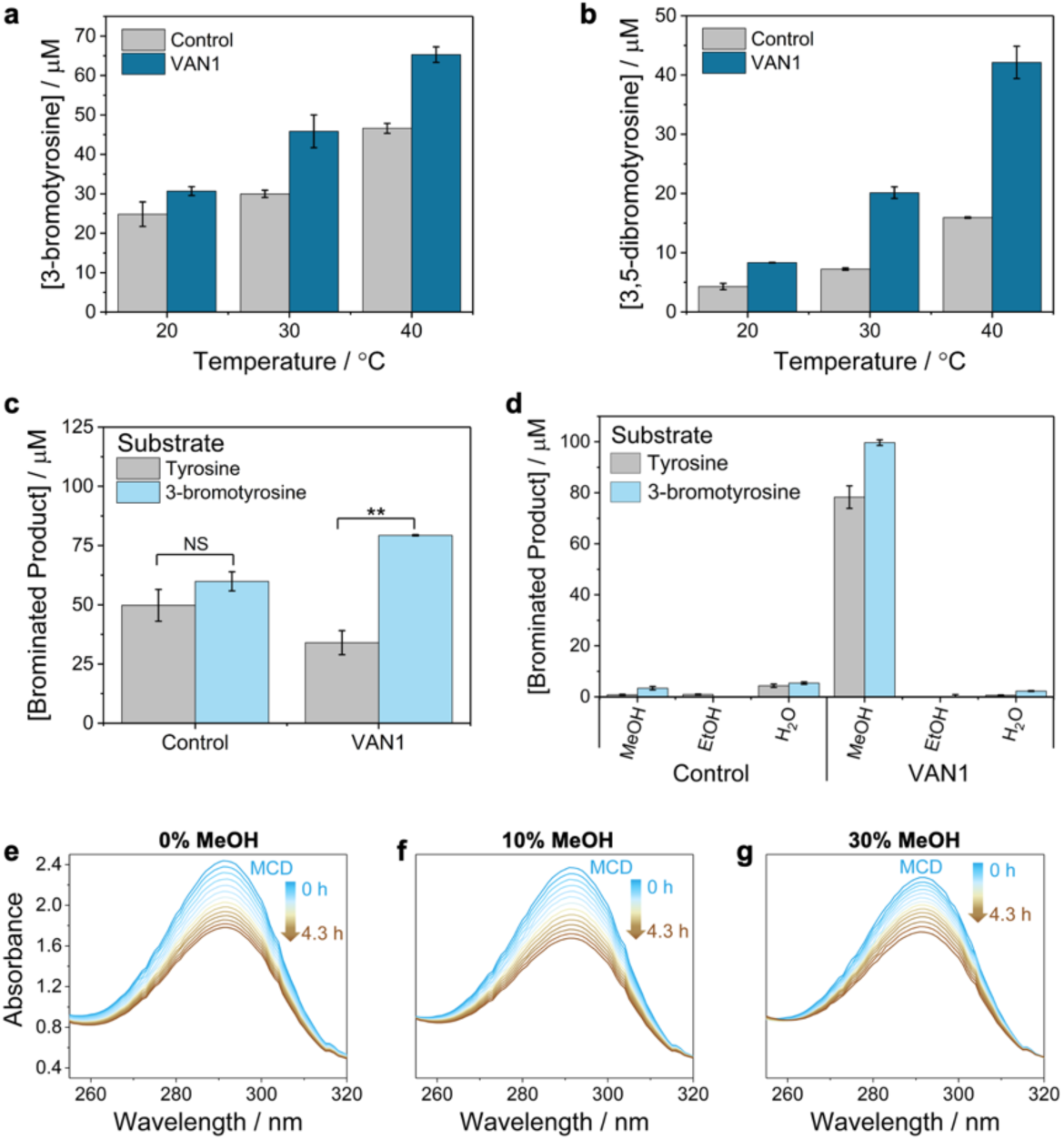
VAN1 and VAN1^E89N^ assays with tyrosine, 3-bromotyrosine, or MCD as the substrate. (a–b) Amount of 3-bromotyrosine (a) or 3,5-dibromotyrosine (b) generated after 6 h under standard aqueous assay conditions at different reaction temperatures with 1 mM *L*-tyrosine as the substrate. Panel b is the same data shown in Fig. 6c and was included for comparison with panel a. (c) Aqueous assays comparing the total brominated product after 6 h using 1 mM tyrosine or 1 mM 3-bromotyrosine as the substrate. Where tyrosine was the substrate, [brominated product] = [3-bromotyrosine] + 2[3,5-dibromotyrosine]; for reactions with 3-bromotyrosine, [brominated product] = [3,5-dibromotyrosine]. Significance thresholds: *p < 0.05, **p < 0.01. (d) Tyrosine bromination assays in which the reactions were diluted 2-fold in either methanol, ethanol, or water to measure enzyme activity in 50% of each solvent. The reaction conditions were identical to those in panel c prior to dilution in each solvent. (e–g) Control MCD kinetic scans at 0–30% methanol. Note: reactions labeled “control” are identical to other conditions but without added protein.

## Supporting Tables

**Table S1.**
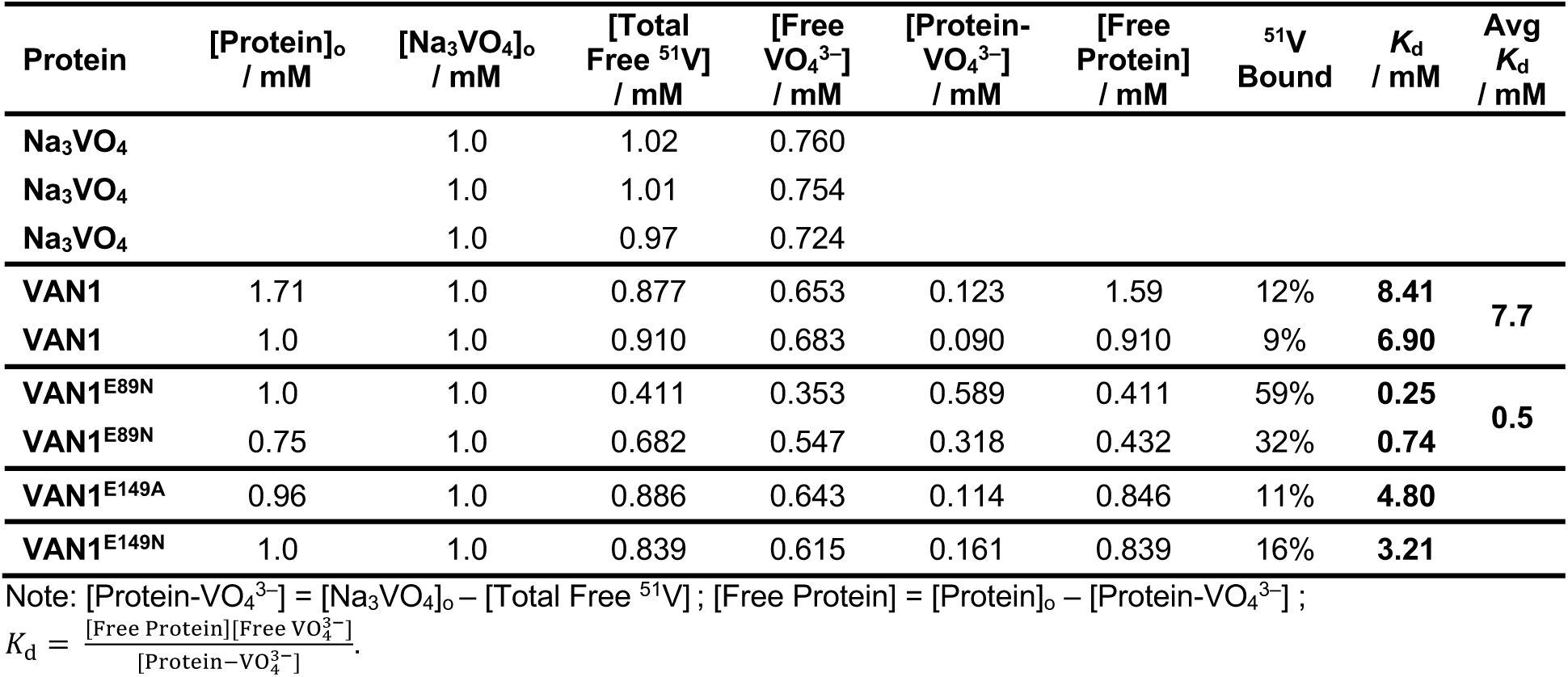
^51^V NMR data used to calculate protein-VO_4_^3–^ binding affinities.

**Table S2.**
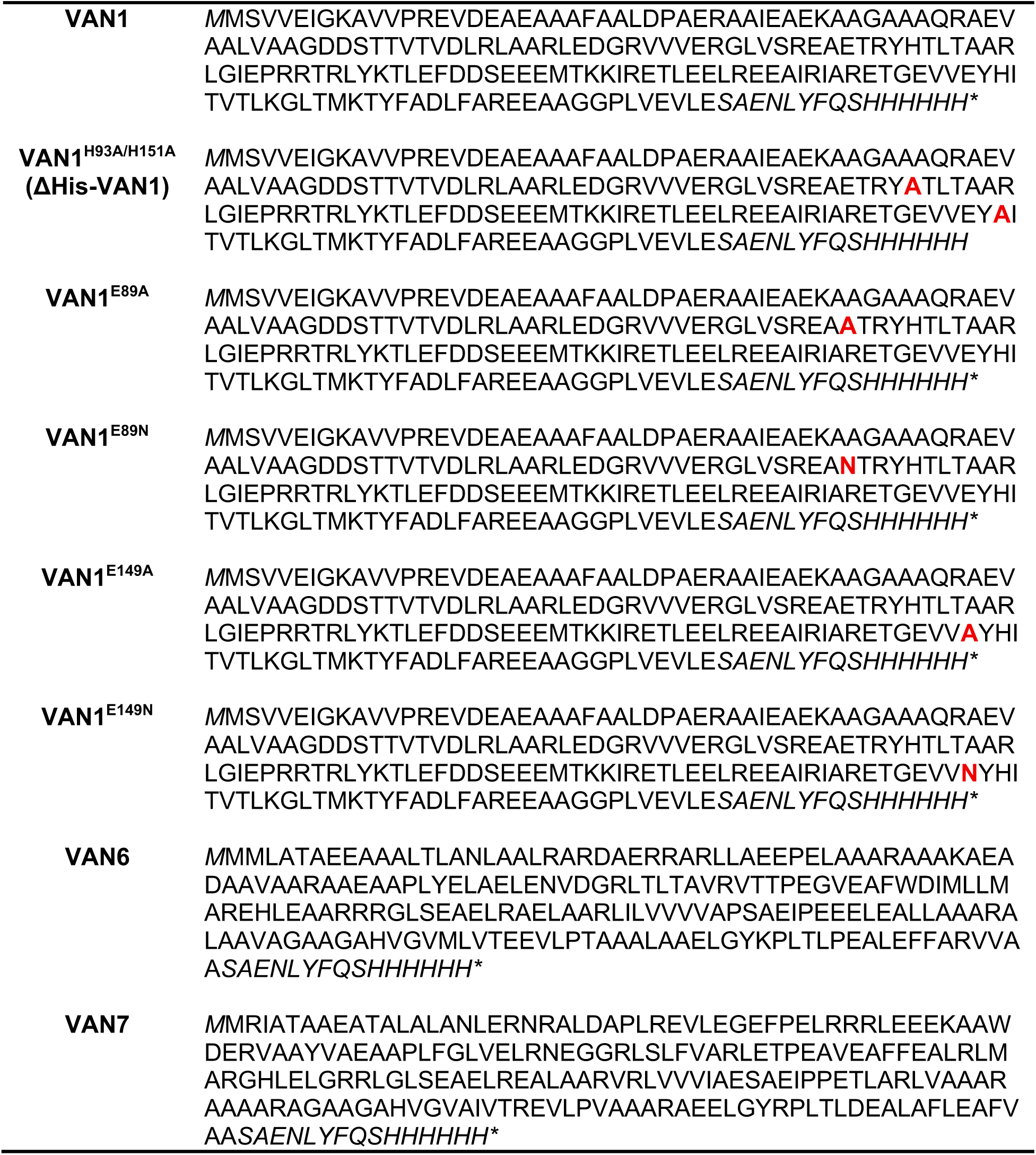
Sequences of proteins used in this work. An added Met at the start and the TEV cleavage site + C-terminal His-tag are italicized. Mutations are shown in bold red text.

**Table S3.**
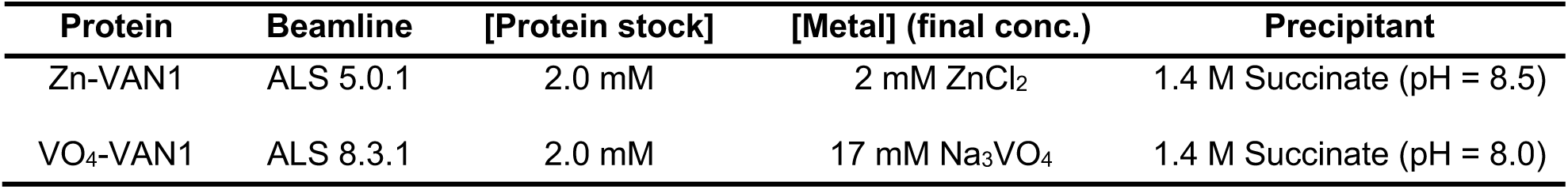
Crystallization conditions for single crystal X-ray diffraction structures reported in this manuscript.

**Table S4.** X-ray data collection and refinement statistics. Numbers in parentheses correspond to the highest resolution shell.

| **Data Collection** | **Zn-VAN1** | **VO<sub>4</sub>-VAN1** |
| --- | --- | --- |
| PDB ID | 39IG | 39JA |
| Space Group | 19 | 19 |
| Unit Cell Dimensions (Å) | 84.09, 84.14, 315.12 | 84.82, 84.80, 319.23 |
| Unit Cell Angles (°) | 90, 90, 90 | 90, 90, 90 |
| Resolution (Å) | 47.47 – 2.69 | 47.94 – 3.44 |
| No. Unique Reflections | 119917 (8892) | 31482 (2331) |
| R<sub>merge</sub> | 0.373 (6.217) | 0.291 (3.158) |
| R<sub>meas</sub> | 0.378 (6.299) | 0.315 (3.439) |
| Multiplicity | 41.8 (38.4) | 6.87 (6.42) |
| CC 1/2 | 0.999 (0.187) | 0.999 (0.134) |
| < I / $\sigma(I)$ > | 10.66 (0.75) | 5.78 (0.64) |
| Completeness (%) | 100.00 (100.00) | 100.00 (100.00) |
| **Refinement** | **Zn-VAN1** | **VO<sub>4</sub>-VAN1** |
| Number of Protein Chains | 8 | 8 |
| Twin Operator | k, h, -l | k, h, -l |
| Estimated Twin Fraction (Xtriage) | 0.478 | 0.478 |
| Refined Twin Fraction (phenix.refine) | 0.500 | 0.500 |
| R<sub>work</sub> / R<sub>free</sub> | 0.308 / 0.308 | 0.365 / 0.372 |
| Non-Hydrogen Atoms | 11102 | 11012 |
| Protein | 11094 | 10987 |
| Ligand/Ion | 8 | 25 |
| Solvent | 0 | 0 |
| RMSD Bonds | 0.021 | 0.007 |
| RMSD Bond Angles | 1.161 | 1.133 |
| Clashscore | 25.31 | 22.95 |
| Ramachandran Favored (%) | 96.91 | 98.21 |
| Ramachandran Outliers (%) | 0.14 | 0.14 |
| Rotamer Outliers (%) | 1.98 | 0.45 |

## Supporting Data

**Supporting Data 1.** RFdiffusion script used to generate VAN1 scaffold.

VAN1:

nohup python RFdiffusion/scripts/run_inference.py inference.num_designs=25

inference.output_prefix=1IDQ_correct_AA_order_version11_VO4Substrate_6_conserved_A As_Outputs/1IDQ_VO4Sub_6_conserved_AAs_RFDiffusion inference.input_pdb=1idq.pdb ’contigmap.contigs=[15-35/A292-292/20-25/A353-353/20-50/A360-360/20-50/A404-404/10-25/A490-490/20-50/A496-496/15-50]’ potentials.guide_scale=3

’potentials.guiding_potentials=["type:substrate_contacts,s:1,r_0:8,rep_r_0:5.0,rep_s:2,rep_r_ min:1"]’ potentials.substrate=VO4

inference.ckpt_override_path=RFdiffusion/models/ActiveSite_ckpt.pt &

**Supporting Data 2.** Shell script used to run ProteinMPNN to obtain the sequences of VAN1– VAN4.

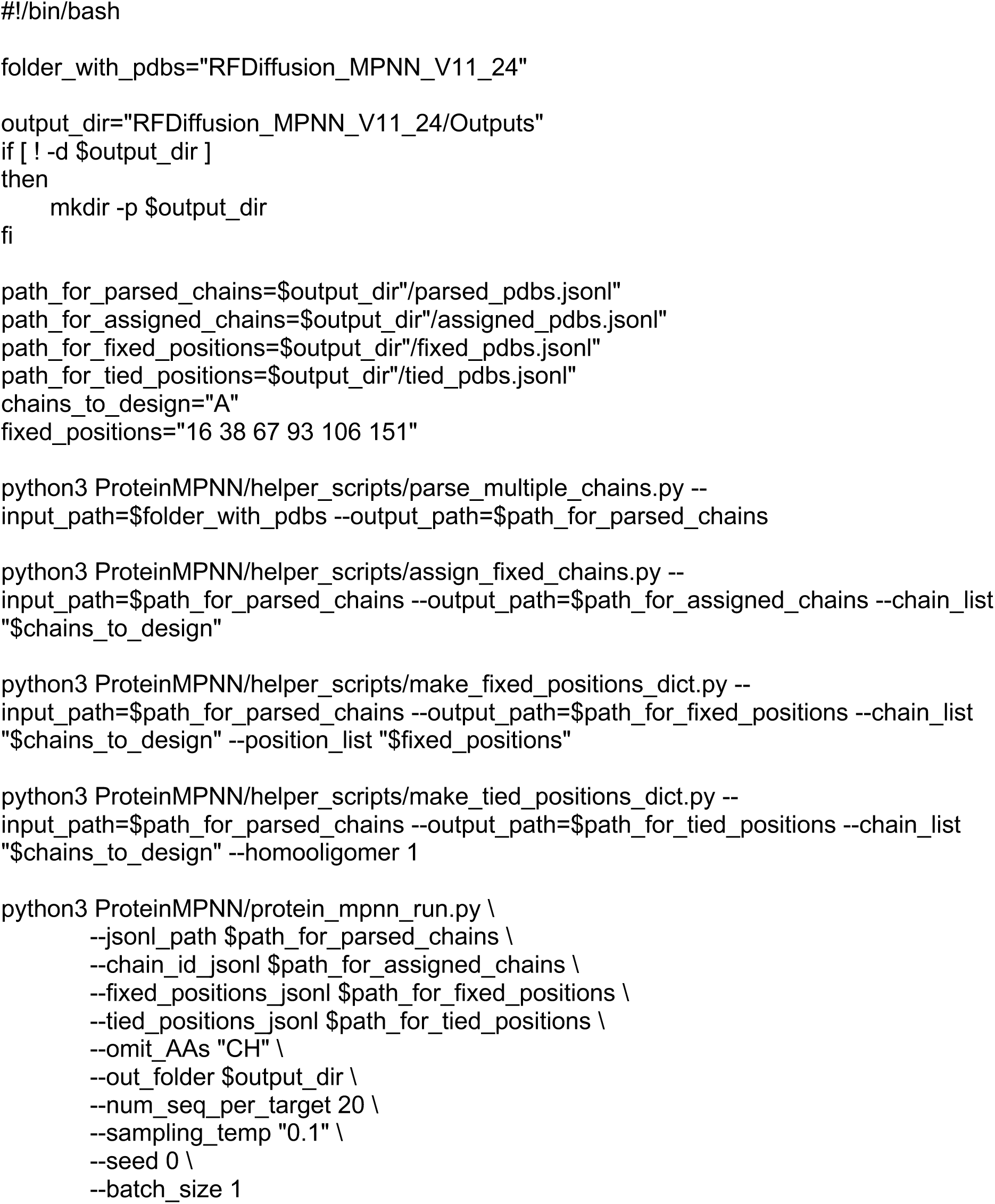

